# Individual differences in decision-making shape how mesolimbic dopamine regulates choice confidence and change-of-mind

**DOI:** 10.1101/2024.09.16.613237

**Authors:** Adrina Kocharian, A. David Redish, Patrick E. Rothwell

**Affiliations:** Graduate Program in Neuroscience, University of Minnesota Medical School, Minneapolis, MN; Medical Scientist Training Program, University of Minnesota Medical School, Minneapolis, MN; Department of Neuroscience, University of Minnesota Medical School, Minneapolis, MN

## Abstract

Nucleus accumbens dopamine signaling is an important neural substrate for decision-making. Dominant theories generally discretize and homogenize decision-making, when it is in fact a continuous process, with evaluation and re-evaluation components that extend beyond simple outcome prediction into consideration of past and future value. Extensive work has examined mesolimbic dopamine in the context of reward prediction error, but major gaps persist in our understanding of how dopamine regulates volitional and self-guided decision-making. Moreover, there is little consideration of individual differences in value processing that may shape how dopamine regulates decision-making. Here, using an economic foraging task in mice, we found that dopamine dynamics in the nucleus accumbens core reflected decision confidence during evaluation of decisions, as well as both past and future value during re-evaluation and change-of-mind. Optogenetic manipulations of mesolimbic dopamine release selectively altered evaluation and re-evaluation of decisions in mice whose dopamine dynamics and behavior reflected future value.

## INTRODUCTION

While making decisions, agents evaluate and re-evaluate paths traveled, comparing to alternate pasts or possible futures^1,2^. These processes are influenced by the confidence with which agents make decisions, as they deliberate the possible future consequences of their choices, or change their minds after considering unchosen options^3–7^. Dopamine in the nucleus accumbens (NAc) core is a key component of the information processing underlying decision-making, implicated in motivation and reward prediction error (RPE) derived from evaluating actual outcomes^8–18^. Dopamine is well-established as a key input to and driver of NAc core function^19–22^, but it remains unknown what information dopamine provides during change-of-mind decisions. Moreover, given that change-of-mind decisions depend on confidence in the original decision, it is also important to investigate how dopamine signals relate to the confidence with which decisions are made.

We monitored and manipulated dopamine dynamics during internally-driven evaluation and re-evaluation processes, using an economic foraging task where mice are known to make both evaluation and re-evaluation (change-of-mind) decisions^19,23–27^. On this task, mice have a limited time budget to spend seeking food rewards of varying subjective value. The self-paced nature of the task allowed for mice to evaluate and re-evaluate decisions by exhibiting change-of-mind behaviors. We found that dopamine tracked decision confidence during evaluation, as well as values from the past and future when an animal changed its mind. However, the nature of these dopamine signals depended upon individual differences in the behavioral strategy used during task performance. In all mice, dopamine levels dipped just before a change-of-mind decision, but optogenetic inhibition of dopamine release selectively increased change-of-mind behaviors in mice sensitive to future value. Mice with this decision-making phenotype also showed increased decision confidence following optogenetic stimulation of dopamine release, and this effect carried over to decrease subsequent change-of-mind behaviors. Together, these findings expand our framework for dopamine in the context of reward prediction error, by elaborating specific conditions where strategy, confidence, and consideration of the future are linked to dopaminergic signaling during decision-making.

## RESULTS

### Assessment of neuroeconomic decision-making with the Restaurant Row task

We trained mice of both sexes^28^ in a sequential foraging task known as “Restaurant Row” that has previously been used to study neuroeconomic decision-making in both mice and rats^19,23–27,29^. In this task, mice have a daily budget of one hour to obtain food rewards of four distinct flavors. As mice ran counterclockwise around the maze, they encountered a different “offer zone” at each corner (Fig. 1a). Upon entering the offer zone, a tone was presented at a frequency that signaled the delay the animal would have to wait until a reward was delivered (Fig. 1b). The delays were random, ranging from 1-30 seconds, and scaled with frequency such that longer delays were signaled by higher-pitched tones. In the offer zone, mice could either skip the offer and continue to the next restaurant, or accept the offer and advance into the “wait zone”. Upon entering the wait zone, tones were presented once per second at decreasing frequencies, indicating the amount of time remaining in the countdown. Mice earned a reward if they waited out the entirety of the delay, but at any point during the countdown, they could quit the trial early by leaving the wait zone for the next restaurant. Quit and skip decisions were followed by quantitatively different behavioral responses to offers at the subsequent restaurant (Extended Data Fig. 1), suggesting they result from distinct decision-making processes.

**Figure 1.**
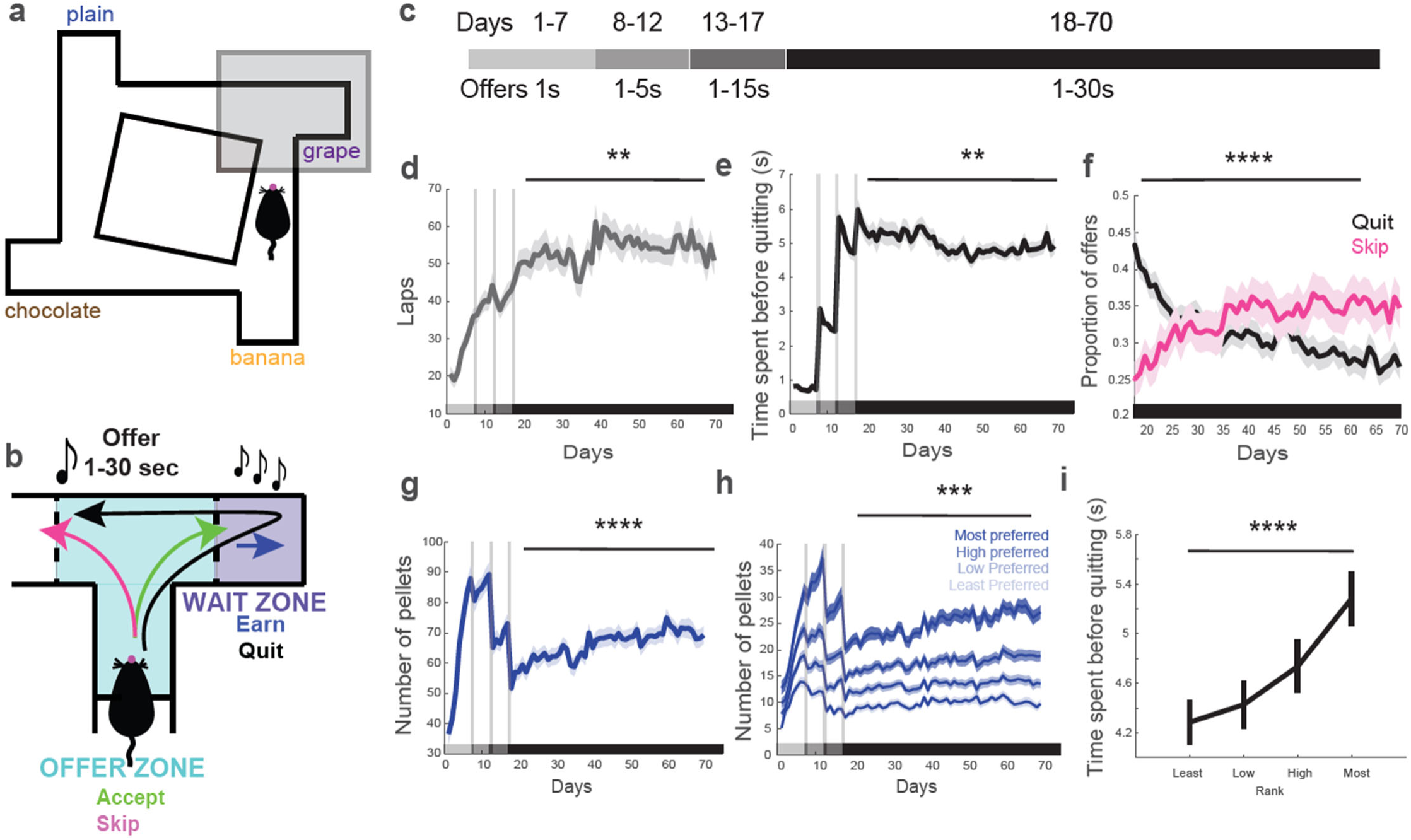
Assessment of neuroeconomic decision-making with the Restaurant Row task. (**a**) Task structure: mice traversed a maze with four restaurants dispensing distinct flavors. (**b**) Single restaurant showing offer zone, wait zone, and potential choices. (**c**) Task training schedule. (**d**) Laps run. (**e**) Time spent before quitting. (**f**) Proportion of quits and skips. (**g**) Pellets earned. (**h**) Earnings by individual restaurant rank. (**i**) Time spent before quitting by rank. Data are mean +/- SEM for all panels. *p<0.05, **p<0.01, ***p<0.001, ****p<0.0001, ANOVA main effect of Day; complete statistics are provided in Supplementary Tables.

Mice were trained on this task over at least 70 days, and were initially presented with short offers that became progressively longer with training (Fig. 1c). When confronted with the full range of offers (1-30 seconds) from day 18 onward, mice developed economic strategies to increase earnings during the limited time available in each behavioral session (one hour). These strategies included increasing the number of laps run (Fig. 1d; F_9.96,436.7_ = 2.39, p = 0.009), decreasing the amount of time invested before quitting (Fig. 1e; F_7.16,313.9_ = 3.32, p = 0.0018), and gradually switching from quitting to skipping behaviors (Fig. 1f; skip: F_8.62,377.8_ = 5.604, p < 0.0001; quit: F_9.49,416.2_ = 10.89, p < 0.0001). This ultimately resulted in a greater number of pellets earned across training from day 18 onward (Fig. 1g; F_10.29,451.1_ = 9.90, p < 0.0001). Individual mice also developed flavor preferences over time (Fig. 1h; main effect of flavor: F_1.42,74.56_ = 151.84, p < 0.0001), and the time spent before quitting scaled up according to flavor preference (Fig. 1i; F_1.94,85.24_ = 49.89, p < 0.0001).

### Individual differences in offer sensitivity define two decision-making phenotypes

As training progressed, we noted a substantial degree of heterogeneity in how individual mice responded to information presented in the offer zone. Some mice showed behavioral sensitivity to tone presentation in the offer zone, more frequently accepting offers with shorter delays and skipping offers with longer delays (Fig. 2a). Other mice accepted short- and long-delay offers with similar probability, exhibiting minimal behavioral sensitivity to offer presentation (Fig. 2b). To quantify offer sensitivity, we calculated the difference in probability of accepting “good” offers (1-3 seconds) and “bad” offers (27-30 seconds) for each individual mouse. The distribution of offer sensitivity included mice close to zero (relatively insensitive) and greater than zero (relatively sensitive). We fit this distribution using Gaussian mixture models with varying numbers of components, and found that the Akaike Information Criterion was minimized by a model with two components (Fig. 2c). We used this two-component Gaussian mixture model to separate “offer-sensitive” mice from “offer-insensitive” mice that exhibited little or no offer sensitivity. Both groups included a similar number of females and males, earned a similar number of pellets, and exhibited similar lingering time (defined as the time between earning a pellet and leaving the wait zone) as a function of flavor (Extended Data Fig. 2a-e).

**Figure 2.**
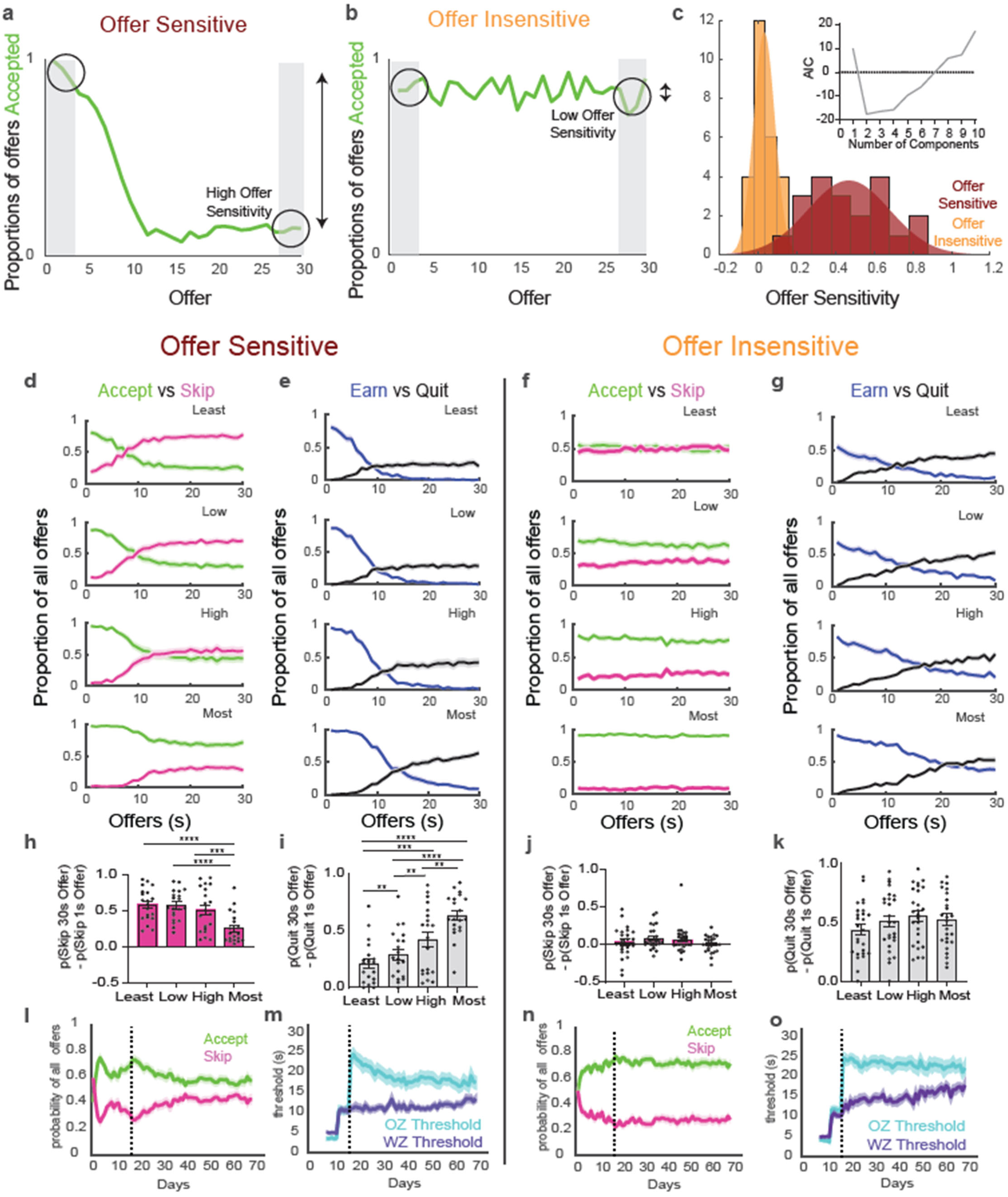
Individual differences in offer sensitivity define two decision-making phenotypes. (**a**) Example of an offer-sensitive mouse whose probability of accepting an offer varies as a function of offer. (**b**) Example of an offer-insensitive mouse whose probability of accepting is relatively consistent across offers. (**c**) Gaussian mixture models with two components fit to the distribution of offer sensitivity, separating the offer-sensitive and offer-insensitive phenotypes. Inset: Akaike information criterion was minimized by a model with two components. (**d to g**) Proportion of offers accepted or skipped by rank and offer for offer-sensitive (d) and offer-insensitive (f), and proportion of offers quit or earned by rank for offer-sensitive (e) and offer-insensitive (g). (**h to k**) Difference in skip probability between high and low offers by rank for offer-sensitive (h) and offer-insensitive (j), and difference in quit probability between high and low offers by rank for offer-sensitive (i) and offer-insensitive (j). (**l to o)** Proportion of offers accepted or skipped across training for offer-sensitive (l) and offer-insensitive (n), and thresholds in offer zone and wait zone for offer-sensitive (m) and offer-insensitive (o). Data are mean +/- SEM for all panels; open and filled circles represent female and male mice, respectively. **p<0.01, ***p<0.001, ****p<0.0001, ANOVA main effect followed by Fisher’s LSD post-hoc test; complete statistics are provided in Supplementary Tables.

In offer-sensitive mice, the degree of offer sensitivity varied as a function of flavor (Fig. 2d). The difference in probability of accepting good versus bad offers was most apparent for less-preferred flavors, and diminished for the most-preferred flavor (Fig. 2h; F_2.42,43.64_ = 17.66, p < 0.0001). In offer-insensitive mice, flavor preference also influenced the overall probability of accepting an offer (Fig. 2f), but there was no difference in the probability of accepting good versus bad offers at any flavor (Fig. 2j). These data suggest that offer-sensitive mice are weighing the future consequences of their decision to accept or skip an offer. This was further evidenced by offer-sensitive mice increasing their skipping behavior (Fig. 2l; F_8.10, 144.9_ = 4.06, p = 0.0002), as they learned to selectively accept offers that matched their willingness to wait (Fig. 2m), with a decrease in offer zone threshold across training. In contrast, offer-insensitive mice maintained constant levels of skipping throughout training (Fig. 2n) and showed no change in offer zone threshold (Fig. 2o), indicating that they continued to accept offers greater than their willingness to wait and were thus less sensitive to the future consequences of their decisions.

After accepting an offer and entering the wait zone, all mice were more likely to re-evaluate their decision and quit while waiting out long delays, regardless of their classification as offer-sensitive (Fig. 2e) or offer-insensitive (Fig. 2g). The difference in probability of quitting long versus short delays also varied as a function of flavor in offer-sensitive mice (Fig. 2i; F_1.92,34.54_ = 30.46, p < 0.0001), but not offer-insensitive mice (Fig. 2k). Time invested before quitting followed a gamma distribution in both groups (Extended Data Fig. 2f). Neither group appeared to quit randomly, neither group appeared to wait for a specific tone in the countdown to cue them to quit, nor did either group appear to wait a fixed number of seconds before quitting (Extended Data Fig. 2g-h).

### NAc core dopamine dynamics reflect decision-making differences

To determine how these decision-making differences were related to mesolimbic dopamine dynamics, we used fiber photometry to monitor dopamine signals in the NAc core. We infused AAV9-CAG-dLight1.3b into the NAc core to express the dLight1.3b fluorescent biosensor^30^ (Fig. 3a), and implanted bilateral fiber optics within the NAc core (Fig. 3b-c and Extended Data Fig. 3). These experiments were conducted using DAT-Cre transgenic mice, so that a second virus could be infused into the VTA to express a Cre-dependent opsin in dopamine neurons and their mesolimbic axon terminals (Fig. 3d-e). In different cohorts of mice, this second virus was either AAVdj-hSyn-DIO-ChrimsonR-tdTomato (expressing a red-shifted excitatory opsin^31^) or AAV5-EF1a-DIO-eNpHR3.0-mCherry (expressing an inhibitory halorhodopsin^32^). This experimental design allowed us to stimulate or inhibit mesolimbic axon terminals in the NAc core with red light, through the same optic fiber used to monitor dLight fluorescence with blue light^33^ (Fig. 3f).

**Figure 3.**
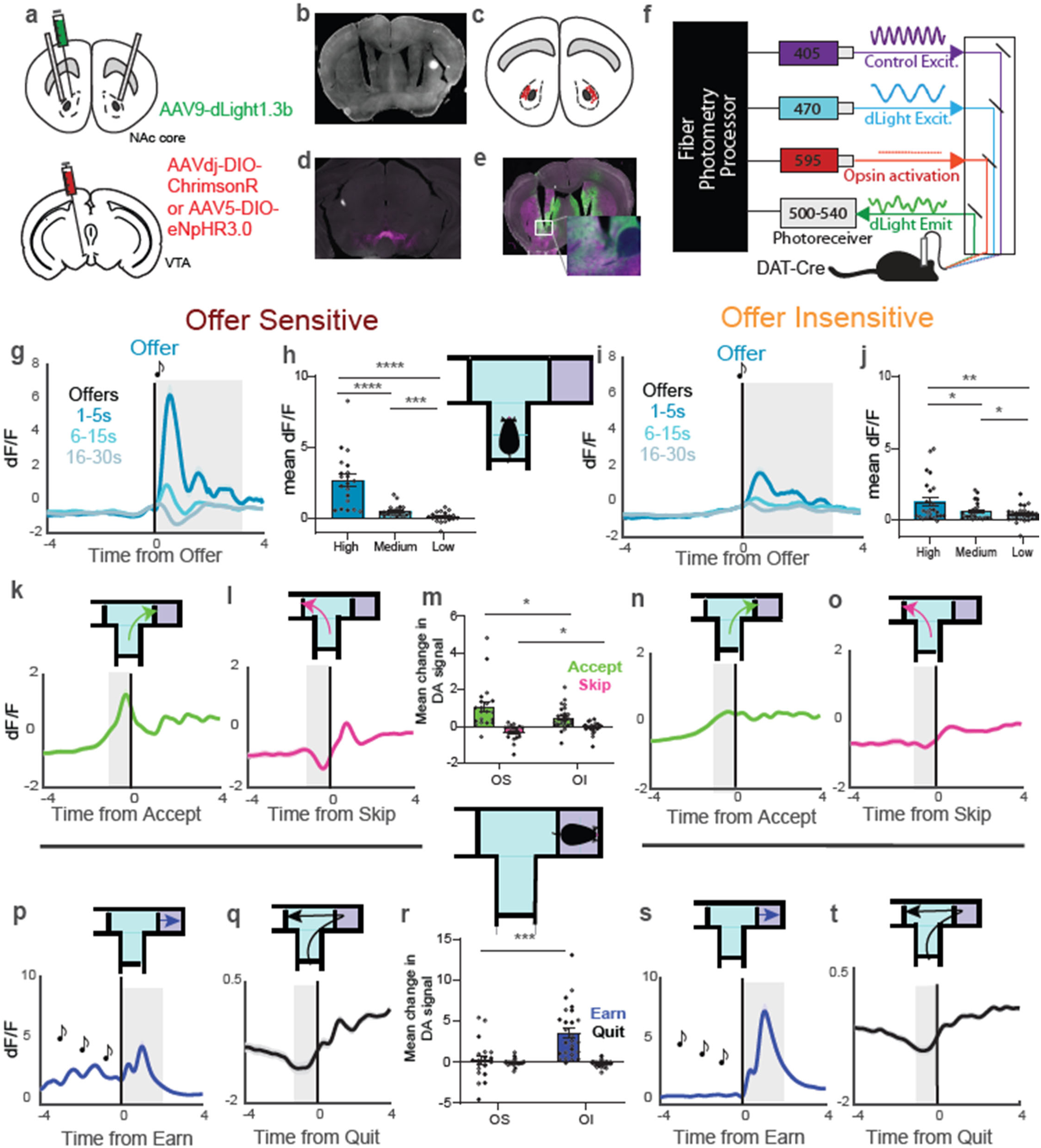
NAc core dopamine dynamics reflect decision-making differences. (**a**) Schematic of virus transfection sites in NAc core (top) and VTA (bottom). (**b**) Example of fiber optic placement. (**c**) Fiber optic tip placements for all animals. (**d**) Example histology showing viral expression of opsin in VTA (magenta). (**e**) ChrimsonR/halorhodopsin terminal expression (magenta) and dLight expression (green) in NAc. (**f**) Fiber photometry recording setup. (**g to j**) Time course of dopamine response to offer in offer-sensitive (g) and offer-insensitive (j), along with average response to high, medium, and low delay offers in offer-sensitive (h) and offer-insensitive (j). (**k to o**) Time course of dopamine responses to accepted offers in offer-sensitive (k) and offer-insensitive (n); skipped offers in offer-sensitive (l) and offer-insensitive (o); and average responses (m). (**p to t**) Time course of dopamine response to earning a pellet in offer-sensitive (p) and offer-insensitive (s); quitting in offer-sensitive (q) and offer-insensitive (t); and average responses (r). Shaded gray boxes indicate time windows used for quantification. Data are mean +/- SEM for all panels; open and filled circles represent female and male mice, respectively. *p<0.05, **p<0.01, ***p<0.001, ****p<0.0001, ANOVA main effect followed by Fisher’s LSD post-hoc test (h and j) or interaction followed by simple effect tests (m and r); complete statistics are provided in Supplementary Tables.

Early in training, offer-sensitive and offer-insensitive mice showed comparable dopamine responses after earning a pellet regardless of flavor, though this dopamine response varied as a function of offer only in offer-sensitive mice (Extended Data Fig. 4a-d). After offer delays reached 1-30 seconds, we compared dopamine signals aligned to the onset of offers with a short delay (1-5 s), medium delay (6-15 s), or long delay (15-30 s). The dopamine response was inversely related to offer length in both offer-sensitive (Fig. 3g-h; F_1.04,18.81_ = 33.38, p < 0.0001) and offer-insensitive mice (Fig. 3i-j; F_1.08,25.90_ = 8.47, p = 0.0063), indicating dopamine signals varied with delay to reward and implying a neurochemical representation of expected cost in both groups. However, the magnitude of this effect was greater in offer-sensitive mice, as indicated by a significant interaction between Offer Length and Offer Sensitivity (F_2,84_ = 11.88, p < 0.0001). The average dopamine signal showed a transient peak once per second in both groups, corresponding to each individual tone presentation during the offer, and providing further evidence that offer-sensitive and offer-insensitive mice could all perceive the offer tone. Importantly, these data indicate that neural signals in offer-insensitive mice reflected offer-based valuation, even if their behavior did not.

We then analyzed dopamine dynamics within the offer zone by separating decisions to accept or skip. Offer-sensitive mice showed bidirectional dopamine responses while evaluating decisions and their future consequences, with dopamine increasing prior to accepting and decreasing prior to skipping an offer (Fig. 3k-m). Offer-insensitive mice also showed a smaller increase in dopamine signal prior to accepting an offer, but no decrease in signal on skip trials (Fig. 3m-o). This pattern was reflected statistically as a significant interaction between wait zone Outcome (accept/skip) and Offer Sensitivity (F_1,42_ = 7.78, p = 0.0002).

After accepting an offer and proceeding into the wait zone, both groups exhibited bidirectional dopamine dynamics based on outcome (earn versus quit). Dopamine tracked the countdown itself in offer-sensitive mice, with small increases in dopamine occurring each second as individual countdown tones were presented. After mice waited through the entire countdown and earned a pellet, we observed increases in dopamine at the time of pellet delivery, which were less robust in offer-sensitive mice (Fig. 3p) and more robust in offer-insensitive mice (Fig. 3s). Conversely, when mice changed their mind and quit the countdown early, there was a decrease in dopamine signal immediately preceding the quit in both offer-sensitive mice (Fig. 3q) and offer-insensitive mice (Fig. 3t). These patterns were reflected statistically as a significant interaction between wait zone Outcome (earn/quit) and Offer Sensitivity (Fig. 3r; F_1,42_ = 16.67, p = 0.0009). Overall, dopamine signals were more robust and dynamic in the offer zone for offer-sensitive mice and in the wait zone for offer-insensitive mice, highlighting a correspondence between decision-making phenotype and dopamine signals in the zone where each mouse made its decision. In fact, these dopamine signals correlated with the offer sensitivity of individual mice for accept, skip, and earn events (Extended Data Fig. 4e-h).

### Dopamine dynamics before and after change-of-mind quits during re-evaluation

The Restaurant Row task provides a unique opportunity to study quitting behavior, which occurs when mice accept an offer and enter the wait zone, but then re-evaluate their decision while waiting and volitionally depart before earning a pellet. Importantly, quitting is an opt-in process that requires a change-of-mind and is distinct from skipping (Extended Data Fig. 1), because quitting occurs after the animal has already accepted the offer and inaction during the countdown defaults to earning a pellet. Furthermore, there is no additional information provided to tell the mouse to quit or alter its expectations: the change-of-mind is internally generated and thus cannot be attributed to simple disappointment arising from a violation of expectations beyond the animal’s control. Furthermore, the dip in dopamine signal that occurs immediately before a quit was unique, in the sense that it was not observed when mice performed a physically identical act of leaving the wait zone after earning and consuming a pellet (Extended Data Fig. 5e-h).

To further examine the information encoded by dopamine signals before and after quits, we quantified dopamine dynamics in relationship to the economic past and future. If quitting is a re-evaluation resulting in a change-of-mind decision, we hypothesized that recovery from this re-evaluation after quitting may produce a concomitant rebound in dopamine. To test this, we examined dopamine dynamics after quitting as a function of past time (i.e., the number of seconds invested in the countdown prior to quitting) and past value (i.e., the difference between the offer received and the animal’s willingness to wait; Fig. 4a). High-value trials are those where the animal has a high propensity to wait (i.e., more preferred flavors) and offer delay is short (i.e., low cost), whereas low willingness to wait and long offer delays constitute low-value trials.

**Figure 4.**
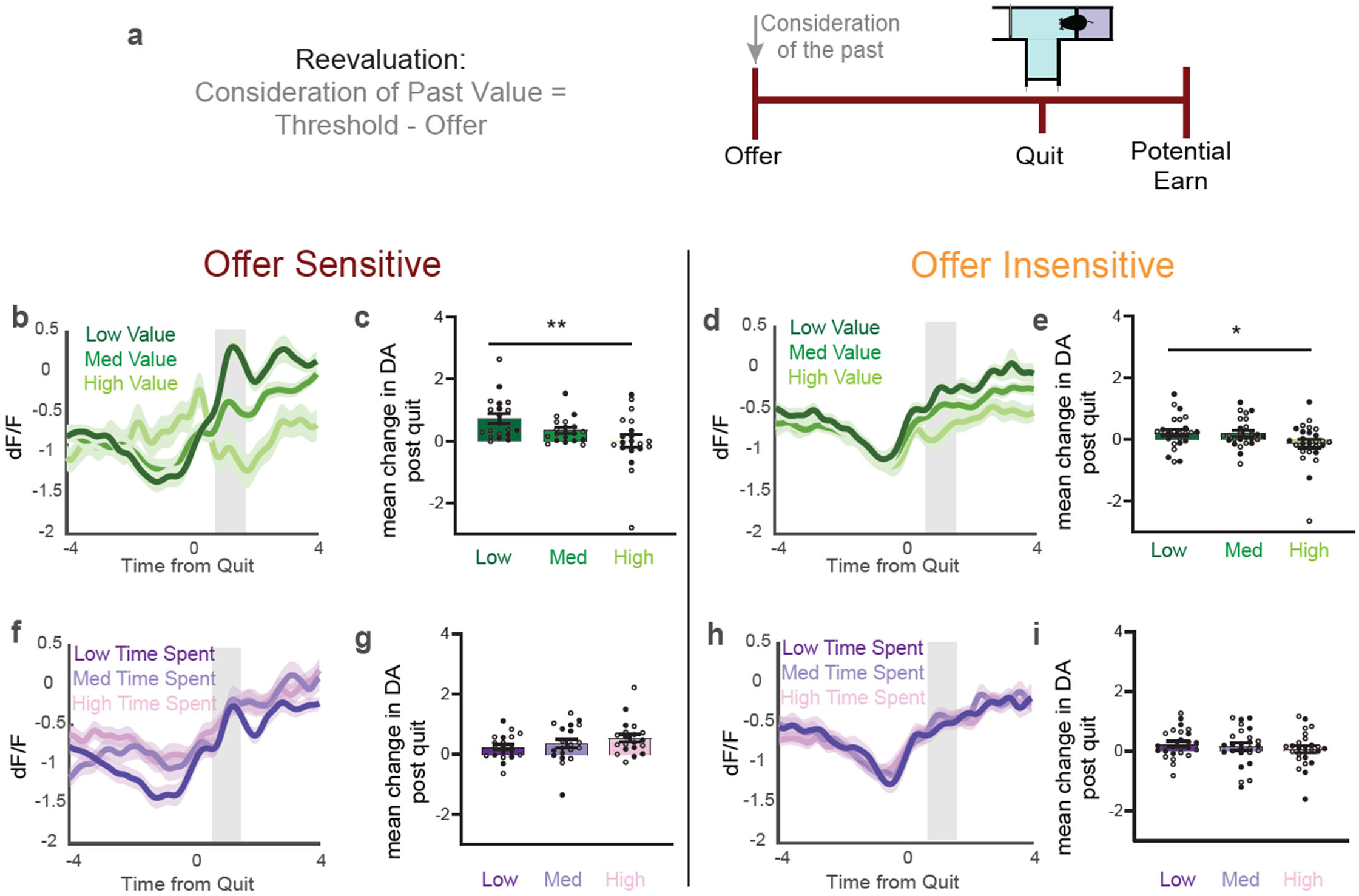
Dopamine rebounds after change-of-mind quitting reflect past value. (**a**) Schematic conceptualizing past value: the difference between willingness to wait (threshold) and offer delay. Highest values refer to short delays at more preferred flavors, while lowest values refer to long delays at less preferred flavors. (**b to e**) Dopamine dynamics during quit rebounds in offer-sensitive (b) and offer-insensitive (e), along with mean change in gray shaded window (c and d). (**f to i**) Dopamine dynamics after quitting with low and high time spent in countdown in offer-sensitive (f) and offer-insensitive (h), along with mean change in gray shaded window (g and i). Data are mean +/- SEM for all panels; open and filled circles represent female and male mice, respectively. *p<0.05, **p<0.01, ANOVA main effect followed by Fisher’s LSD post-hoc test; complete statistics are provided in Supplementary Tables.

After quitting, rebounds in dopamine scaled inversely with past value in both offer-sensitive (Fig. 4b-c; F_2,36_ = 6.45, p = 0.004) and offer-insensitive mice (Fig. 4d-e; F_2,48_ = 3.32, p = 0.045). The magnitude of this effect was similar in both groups, as there was no statistical interaction between Past Value and Offer Sensitivity (F_2,84_ = 1.54, p = 0.22). Low-value trials elicited the largest rebounds in dopamine, whereas high-value trials elicited the smallest rebounds in dopamine. This suggests that the rebound in dopamine signal after quitting reflects consideration of past value. This effect was not seen when animals skipped an offer (Extended Data Fig. 5a-d), or when separating quit trials by past time (Fig. 4f-i), implying the rebound in dopamine signal after quits selectively reflected the re-evaluated and rejected (past) value.

We next analyzed dopamine dynamics during change-of-mind quits in the context of future value. On Restaurant Row, the future can be characterized concretely as the time remaining in the countdown after quitting, or more abstractly as the future value remaining in the countdown after quitting. We can calculate future value by subtracting the time remaining in the countdown at the time of quit from the animal’s willingness to wait (threshold; Fig. 5a). Economically favorable quits occur when the time remaining in the countdown exceeds the animal’s threshold (value remaining < 0), because the animal is relinquishing a future that requires waiting longer than it is typically willing to wait. Thus, favorable quits exemplify consideration of the future value of an imagined prospective reward. Conversely, economically unfavorable quits occur when the time remaining in the countdown is less than the animal’s threshold for that reward (value remaining > 0), because the animal would normally have been willing to wait out the remaining time. Interestingly, we found that offer-sensitive mice had higher rates of favorable quits that increased across days during training (Fig. 5b), while offer-insensitive mice had higher rates of unfavorable quits which remained relatively constant throughout training (Fig. 5c). This supports the notion that the future consequences of decisions had a greater influence on the behavior of offer-sensitive mice.

**Figure 5.**
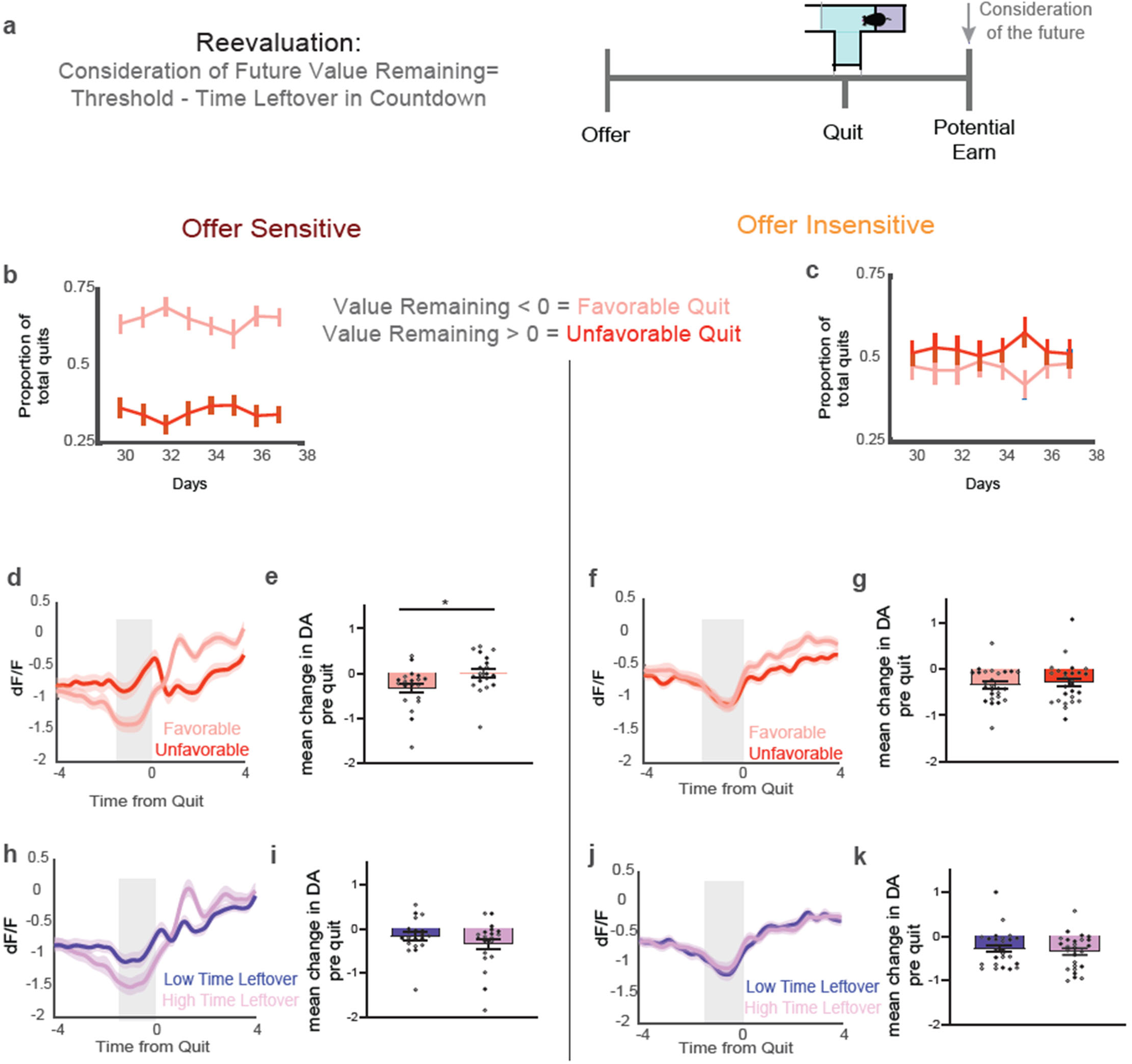
Dopamine dips before change-of-mind quitting reflect future value. (**a**) Schematic conceptualizing future value: the difference between willingness to wait (threshold) and time remaining in the countdown at quit. Values greater than zero imply that it would be economically unfavorable to quit, while values less than zero imply that it would be economically favorable to quit. (**b to c**) Favorable and unfavorable quits in offer-sensitive (b) and offer-insensitive (c). (**d to g**) Dopamine dynamics during favorable and unfavorable quits in offer-sensitive (d) and offer-insensitive (f), along with mean change in gray shaded window (e and g). (**h to k**) Dopamine dynamics while quitting with low and high time remaining in countdown in offer-sensitive (h) and offer-insensitive (j), along with mean change in gray shaded window (i and k). Data are mean +/- SEM for all panels; open and filled circles represent female and male mice, respectively. *p<0.05, ANOVA interaction followed by simple effect test; complete statistics are provided in Supplementary Tables.

Dopamine dynamics at the time of quitting also differed between offer-sensitive and offer-insensitive mice. Offer-sensitive mice showed larger dips in dopamine for favorable versus unfavorable quits (Fig. 5d-e; F_1,18_ = 18.70, p = 0.0004), while offer-insensitive mice showed similar dips in dopamine regardless of quit favorability (Fig. 5f-g). This pattern was reflected statistically as a significant interaction between Quit Favorability and Offer Sensitivity (F_1,42_ = 4.31, p = 0.044). Importantly, these quit-related dips in dopamine signal were not related to future time remaining in the countdown in either offer-sensitive or offer-insensitive mice (Fig. 5h-k). Furthermore, when we controlled for time remaining until earning a pellet, we found the dopamine signal was significantly higher during the countdown for high-value offers than low-value offers (Extended Data Fig. 6), providing further evidence that dopamine signals tracked value more closely than time. In total, these data show that the dip in dopamine signal before a change-of-mind quit was specific to future value in offer-sensitive mice, whereas the rebound in dopamine signal after a change-of-mind quit were specific to past value in both groups. These results provide further evidence that offer-sensitive mice considered the future in a distinct way that differed from offer-insensitive mice.

### Optogenetic inhibition of dopamine release caused change-of-mind quitting in a selective manner

To further test the relationship between dips in dopamine signal and change-of-mind quitting, we used a subset of DAT-Cre transgenic mice that received bilateral injection of Cre-dependent halorhodopsin in the VTA (Fig. 3a). This experimental design allowed us to inhibit mesolimbic dopamine terminals in the NAc core with red light, through the same optic fiber used to monitor dLight fluorescence with blue light (Fig. 6a). We first validated the efficacy of this optogenetic manipulation on dopamine dynamics by measuring dLight signal after delivery of a food pellet, which normally produces a robust increase in dopamine signal (Fig. 6b). To activate halorhodopsin, we delivered light (589 nm, 6-8 mW) for two seconds following pellet delivery (Fig. 6c), which had the expected impact of robustly reducing the dLight signal compared to trials with no light delivery (Fig. 6d; F_1,15_ = 8.24, p = 0.012).

**Figure 6.**
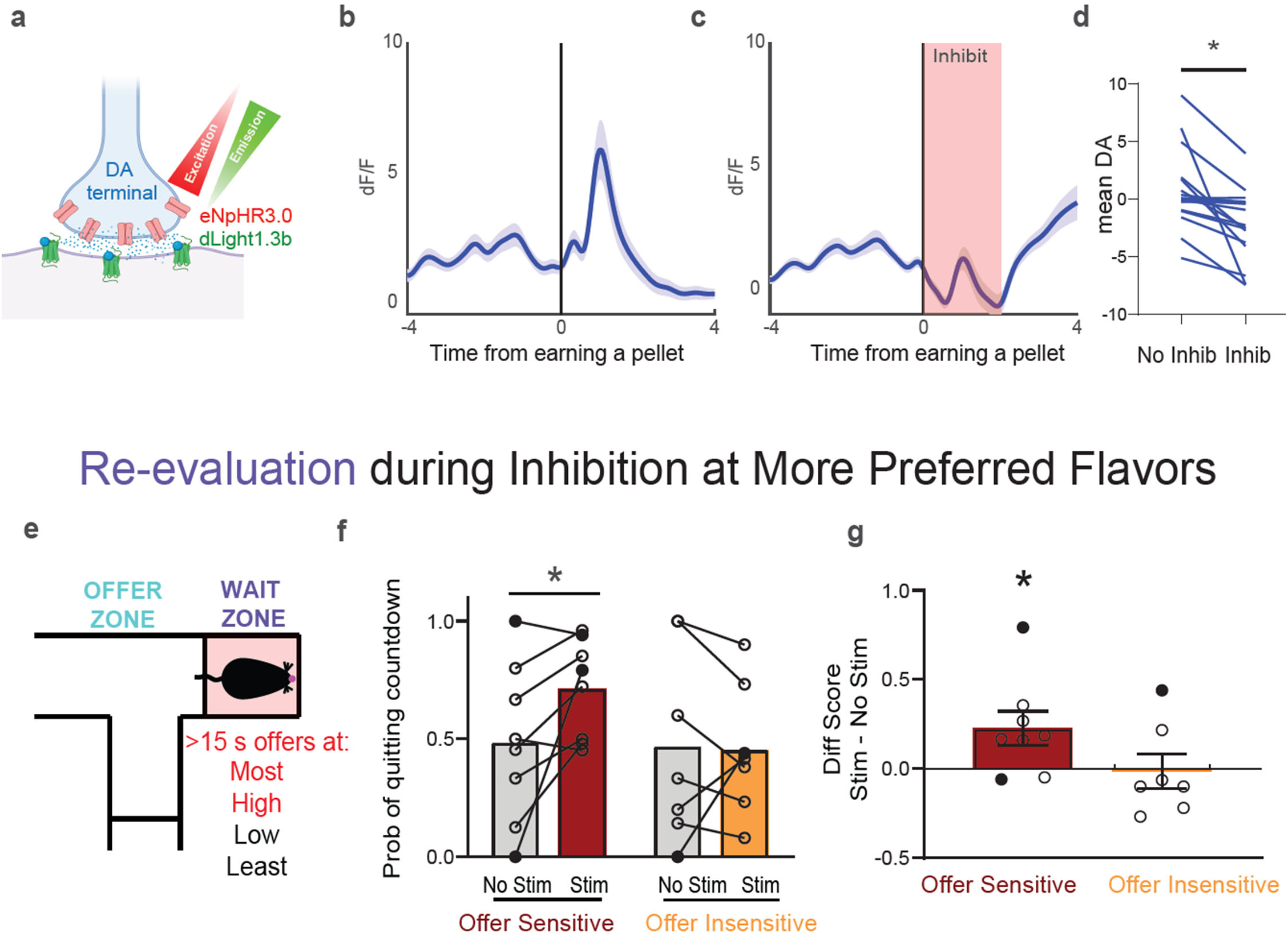
Optogenetic inhibition of dopamine release causes change-of-mind quitting in a selective manner. (**a**) Schematic showing halorhodopsin expression in dopamine terminals, and dLight1.3b expression in the NAc core. (**b to d**) Dopamine response to earning a pellet with (c) or without (b) light delivery, along with average dopamine response (d). (**e**) Wait zone locations (red shading and text) where light was delivered on half of offers > 15 s. (**f**) Probability of quitting on trials without light delivery (No Stim) and with light delivery (Stim) at more preferred flavors. (**g**) Change in probability of quitting caused by light delivery at more preferred flavors. Data are mean +/- SEM for all panels; open and filled circles represent female and male mice, respectively. *p<0.05, simple effect of light delivery in offer-sensitive mice; complete statistics are provided in Supplementary Tables.

This same cohort of mice was trained on the Restaurant Row task, in order to examine how optogenetic inhibition of dopamine release affected behavior. We delivered light on half of all offers >15 sec (Fig. 6e), while the other half of these trials served as an internal control condition with no optogenetic inhibition. Light was initially delivered for only the two more preferred flavors, where probability of quitting is normally low. After mice accepted an offer for a preferred flavor, entry into the wait zone triggered bilateral light delivery for a duration of four continuous seconds, unless they quit and left the wait zone (which ended light delivery). Importantly, the average duration of light delivery was similar in offer-sensitive and offer-insensitive mice (Extended Data Fig. 7a).

Compared to control trials of the same type during the same session but with no light delivery (Fig. 6f), optogenetic inhibition of dopamine release caused a higher probability of quitting in offer-sensitive mice (F_1,7_ = 5.76, p = 0.048), but not offer-insensitive mice (F_1,6_ < 1). This pattern was reflected by a trend toward a statistical interaction between Light Delivery and Offer Sensitivity (F_1,13_ = 3.17, p = 0.098), which is also evident in a comparison of the difference scores (light - control) between groups^34^ (Fig. 6g). We did not observe an effect in offer-sensitive mice when light was delivered for all flavors (Extended Data Fig. 7b-c), ruling out the possibility that optogenetic inhibition of mesolimbic dopamine release caused generic changes in movement. The specific effect of optogenetic inhibition in offer-sensitive mice suggests that individual differences in consideration of the future may influence the way dips in dopamine signal underlie change-of-mind decisions.

### Dopamine dynamics in offer-sensitive mice relate to decision confidence during evaluation

On the Restaurant Row task, consideration of the future also occurs during evaluation of decisions in the offer zone, where mice exhibit reorientation behaviors referred to as vicarious trial and error (VTE). VTE is a well-established behavior in rodents and correlates with future planning and deliberation^35^. During VTE in rodents, neural representations of possible future outcomes sweep serially along different potential paths and alternate between goals, suggesting that animals are considering potential options^36–39^. VTE is best captured by calculating the integrated angular velocity (IdPhi) within the offer zone, which relates to the curvature of the path the animal takes to either accept or skip an offer. We defined VTE events as those represented by high IdPhi values (zIdPhi > 0), which exemplify more variable paths with greater tortuosity (Fig. 7a), and correlate with higher degrees of deliberation. As expected^19^, mice exhibit more VTE prior to a skip decision at more preferred flavors, as offers that are harder to turn down may elicit more deliberation (Extended Data Fig. 8f-g).

**Figure 7.**
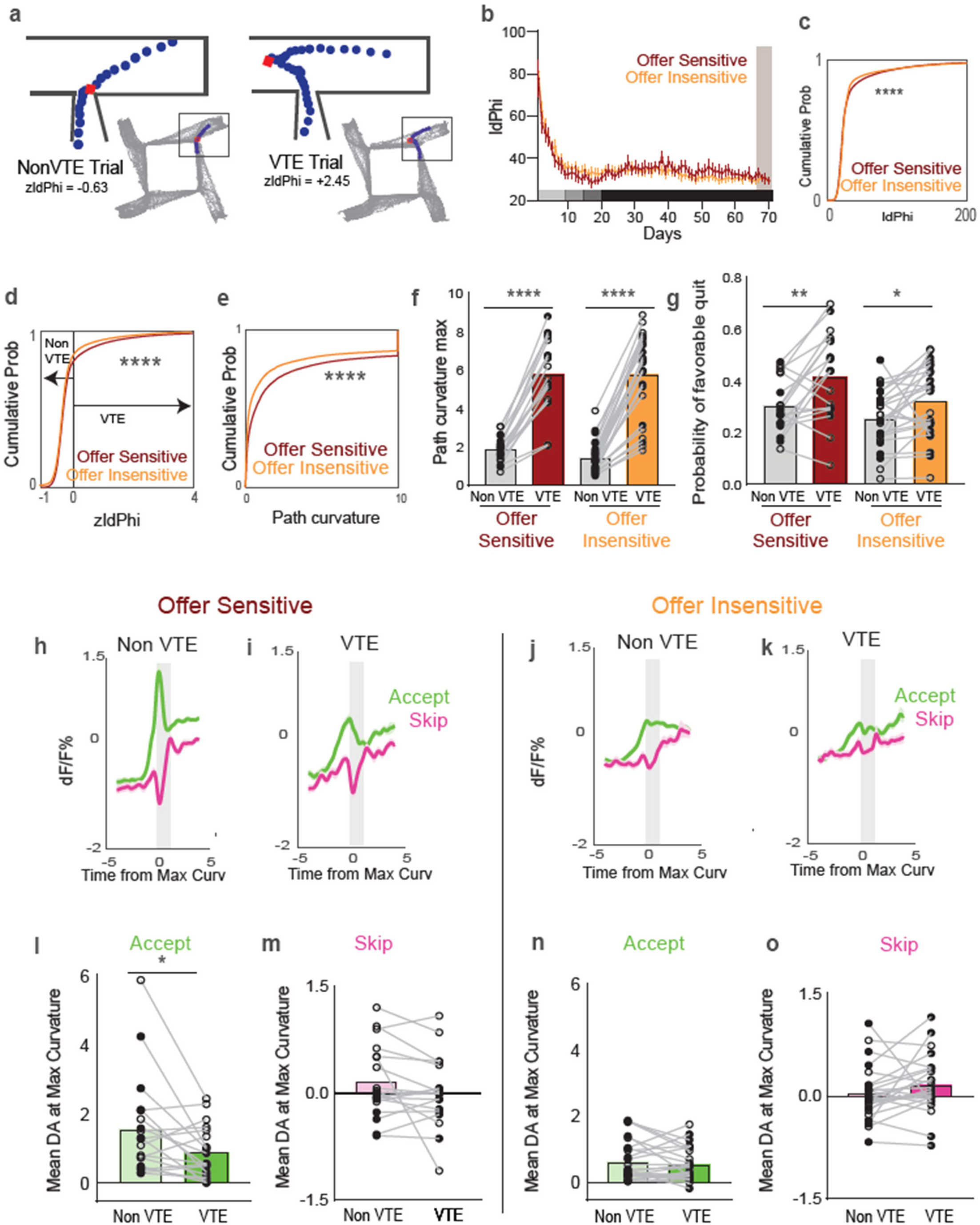
Dopamine dynamics in offer-sensitive mice relate to decision confidence. (**a**) Example path curvatures on trials with low zIdPhi (Non-VTE, left) and high zIdPhi (VTE, right); red square represents point of maximum curvature. (**b**) IdPhi across training; gray shaded area represents dopamine recording days. (**c to e**) Distribution of IdPhi (c) and zIdPhi (d) and path curvatures (e). (**f to g**) Maximum path curvature (f) and proportion on favorable quits (g) on Non-VTE and VTE trials. (**h to j)** Dopamine dynamics aligned to point of maximum path curvature in offer-sensitive on Non-VTE (h) and VTE (i) trials, and offer-insensitive on Non-VTE (j) and VTE (k) trials. (**l to o)** Mean dopamine response in gray shaded window for Non-VTE and VTE trials that were accepted (l) or skipped (m) in offer-sensitive, and accepted (n) or skipped (o) in offer-insensitive. Data are mean +/- SEM for all panels; open and filled circles represent female and male mice, respectively. *p<0.05, **p<0.01, ****p<0.0001, Kolmogorov-Smirnov test (c to e) or ANOVA simple effect (f, g, and l); complete statistics are provided in Supplementary Tables.

As training progressed, offer-sensitive mice began to demonstrate more VTE during evaluation than offer-insensitive mice (Fig. 7b). This was indicated by a rightward shift in the distribution of IdPhi (Fig. 7c), a change in the distribution of VTE and non-VTE trials (Fig. 7d), and a rightward shift in the distribution of max curvature (Fig. 7e). Together, these results suggest that offer-sensitive mice engaged in more deliberation during evaluation, as they considered future consequences of their decisions. As expected, path curvature was greater on VTE trials than non-VTE trials in both groups (Fig. 7f; main effect of VTE: F_1,41_ = 311.6, p < 0.0001). All mice were more likely to make favorable quits after accepting offers on VTE trials than on non-VTE trials (Fig. 7g; main effect of VTE: F_1,41_ = 21.86, p < 0.0001). This suggests that decisions were made with a lower degree of confidence on VTE trials, allowing for quicker re-evaluation and change-of-mind correction, while value remaining was still low.

We next determined the extent to which VTE influenced dopamine dynamics during evaluation in the offer zone. After alignment to the time of peak path curvature to capture deliberative events (Fig. 7a, red squares), we examined dopamine dynamics as mice made decisions to accept or skip offers, separating trials based on the presence or absence of VTE (Fig. 7h-k). Interestingly, when offer-sensitive mice exhibited VTE before accepting an offer, we observed a smaller peak signal compared to accepted offers without VTE (Fig. 7l). Thus, as offer-sensitive mice evaluate the future consequences of their decisions, dopamine levels positively correlated with decision confidence. In contrast, offer-insensitive mice showed comparable dopamine levels on trials with or without VTE (Fig. 7n). This pattern was reflected statistically as a significant interaction between VTE and Offer Sensitivity (F_1,41_ = 6.69, p = 0.0013). In both offer-sensitive and offer-insensitive mice, dopamine dynamics were similar following decisions to skip on trials with or without VTE (Fig. 7, m and o). While these analyses were conducted on data collected after extended training (Fig. 7b, shaded area), similar patterns in dopamine dynamics were also apparent earlier in training (Extended Data Fig. 8a-e).

### Optogenetic enhancement of dopamine release influenced evaluation and re-evaluation of decisions in a selective manner

Since our previous analyses revealed a correlation between dopamine dynamics and decision confidence in offer-sensitive mice, we next used optogenetic stimulation to manipulate dopamine dynamics and measure decision confidence. We used a subset of DAT-Cre transgenic mice that received bilateral injection of Cre-dependent ChrimsonR in the VTA (Fig. 3a). This experimental design allowed us to stimulate mesolimbic dopamine terminals in the NAc core with red light (589 nm, 20 Hz), through the same optic fiber used to monitor dLight fluorescence with blue light (Fig. 8a). Since ChrimsonR expression varied between animals, we leveraged expression of dLight in the NAc core to construct stimulation-response curves for each individual animal. We then tailored optogenetic stimulation parameters for each animal to standardize the evoked signal across mice and to produce responses in a range ∼3-10% dF/F, similar to endogenous signals^40–42^ for offer presentations and pellet consumption (Fig. 8b-c). The average number of pulses selected for each individual animal did not differ between offer-sensitive and offer insensitive-mice (Extended Data Fig. 9c-d).

**Figure 8.**
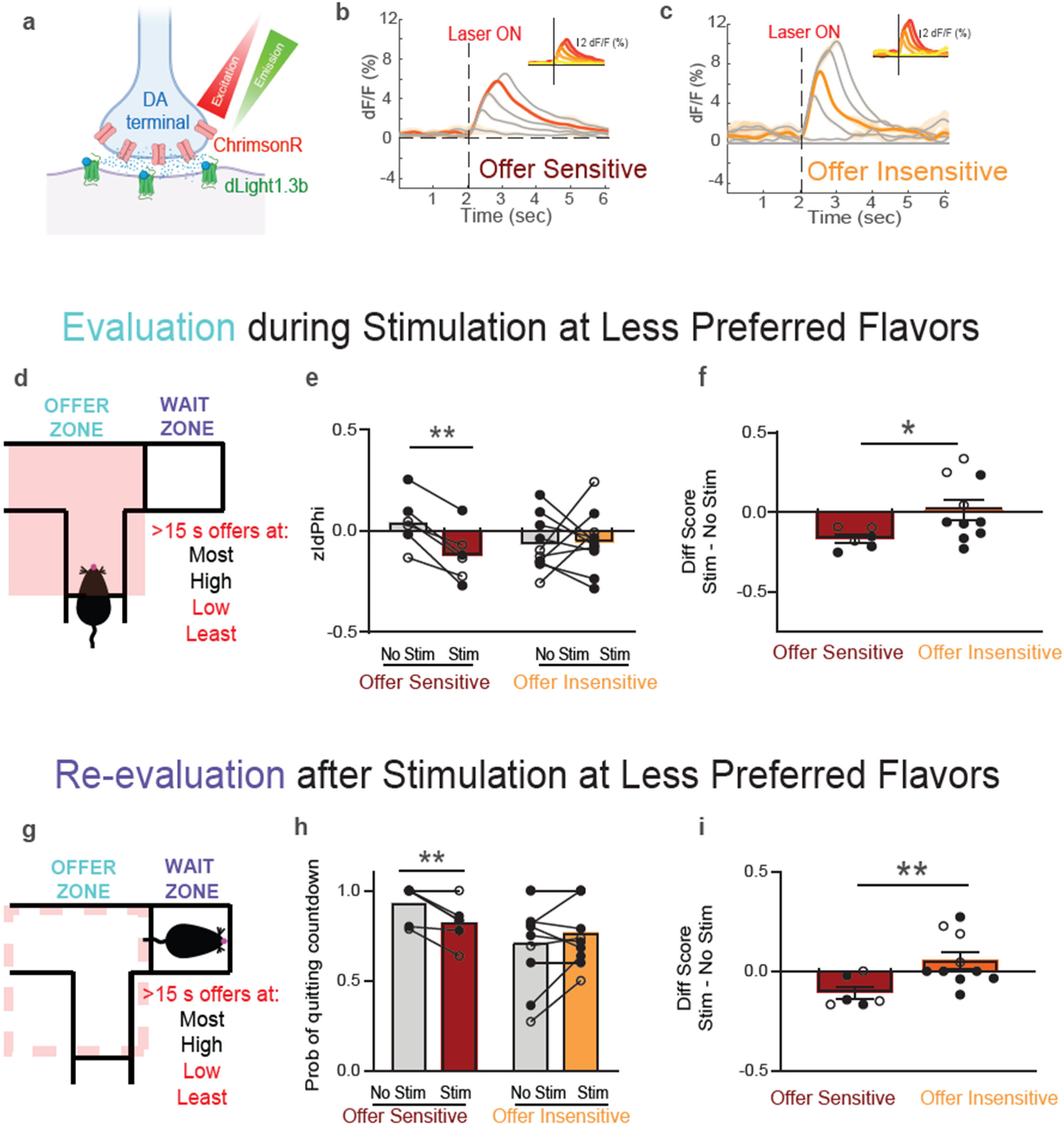
Optogenetic enhancement of dopamine release influences evaluation and re-evaluation of decisions in a selective manner. (**a**) Schematic showing ChrimsonR expression in dopamine terminals, and dLight1.3b expression in the NAc core. (**b to c**) Dopamine response to different pulse numbers (0, 5, 10, 15, or 20; 589 nm, 20 Hz) in offer-sensitive (b) and offer-insensitive (c), highlighting calibrated stimulation parameters. (**d**) Offer zone locations (red shading and text) where optogenetic stimulation was delivered on half of offers > 15 s. (**e**) zIdPhi on trials without (No Stim) and with light delivery (Stim) at less preferred flavors. (**f**) Change in zIdPhi caused by light delivery at less preferred flavors. (**g**) Schematic showing behavior in the wait zone after light delivery in the offer zone (dotted red line) had ended. (**h**) Probability of quitting on trials after no light delivery (No Stim) or light delivery (Stim) at less preferred flavors. (**i**) Change in probability of quitting caused by light delivery at less preferred flavors. Data are mean +/- SEM for all panels; open and filled circles represent female and male mice, respectively. *p<0.05, **p<0.01, simple effect of light delivery (e and h) or offer sensitivity (f and i); complete statistics are provided in Supplementary Tables.

We delivered optogenetic stimulation of dopamine terminals in the offer zone on half of all offers >15 sec (Fig. 8d). Stimulation was initially delivered for only the two less preferred flavors, where probability of offer acceptance is normally low. Compared to control trials of the same type during the same session with no light delivery (Fig. 8e), this manipulation reduced deliberative behaviors (zIdPhi) in offer-sensitive mice (F_1,5_ = 39.19, p = 0.0015), but not offer-insensitive mice. This pattern was reflected statistically as a significant interaction between Light Delivery and Offer Sensitivity (F_1,12_ = 12.47, p = 0.0041), which is also evident in a comparison of the difference scores (light - control) between groups^34^ (Fig. 8f). On separate days of testing, we repeated this experiment but delivered stimulation on half of all offers >15 sec for either the two more preferred flavors or all flavors. These manipulations did not have significant effects on offer-sensitive mice (Extended Data Fig. 10a-d), ruling out the possibility that optogenetic stimulation of mesolimbic dopamine release caused generic changes in movement. Pulsed light delivery had no effect on zIdPhi in offer-sensitive control mice lacking opsin expression (Extended Data Fig. 10i-j), or when delivered in a mismatched fashion to mice expressing halorhodopsin (Extended Data Fig. 7d-i), arguing against the possibility that light delivery was generally distracting. The lack of effect in offer-insensitive mice under identical stimulation conditions suggests that stimulation of dopamine release alters decision-making in a selective manner, possibly by changing the deliberative process by which offer-sensitive mice evaluate the future consequences of accepting an offer.

We also analyzed behavioral responses after mice accepted an offer accompanied by optogenetic stimulation and entered the wait zone (Fig. 8g). Even though light delivery ended when mice entered the wait zone, offer-sensitive mice still showed a decrease in the probability of quitting after prior stimulation at the two less preferred flavors (Fig. 8h; F_1,5_ = 12.13, p = 0.0176), while there was no effect for offer-insensitive mice. This pattern was reflected statistically as a significant interaction between Light Delivery and Offer Sensitivity (F_1,12_ = 12.47, p = 0.0041), which is also evident in a comparison of the difference scores (light - control) between groups^34^ (Fig. 8i). No changes in the probability of quitting were observed in control mice lacking opsin expression (Extended Data Fig. 10k-l), with mismatched light delivery to mice expressing halorhodopsin (Extended Data Fig. 7j), or with stimulation at either the two more preferred flavors or all flavors (Extended Data Fig. 10e-h). The persistent effect of offer zone stimulation during re-evaluation in the wait zone, even after light delivery had ended, suggests that enhanced release of dopamine altered both current and future decision-making for offer-sensitive mice. Importantly, the absence of these effects in offer-insensitive mice suggest that individual differences in decision-making phenotype can dictate the manner in which dopamine dynamics regulate behavioral outcome.

## DISCUSSION

Even when different behavioral strategies lead to the same outcome, the process underlying those decisions may diverge. We found that the process by which mice arrived at decisions was critical in shaping dopamine dynamics: individual differences in decision strategy and dopamine dynamics influenced the extent to which mice expressed confidence in their options and engaged change-of-mind processes to re-evaluate decisions. We identified two behavioral strategies with distinct relationships between dopamine and decision-making. Offer-sensitive mice engaged in more deliberative, future thinking strategies: skipping more economically unfavorable offers during evaluation (in the offer zone) and committing more economically favorable quits during re-evaluation (in the wait zone).

During re-evaluation, we observed dips in dopamine just before animals exhibited change-of-mind behaviors, suggesting that the change-of-mind behaviors were in response to a negative re-evaluation of the situation. Supporting this hypothesis, we found that optogenetic inhibition of dopamine signals increased those change-of-mind behaviors. Further, we discovered that dopamine dips were specific to quitting in economically favorable conditions in offer-sensitive mice, and that dopamine inhibition also specifically increased the probability of quitting the countdown in offer-sensitive mice. This suggests that dopamine dips may causally reflect a cognitive re-evaluation process that underlies change-of-mind specifically in animals who are more future-thinking.

In line with this hypothesis, dopamine dynamics also reflected decision confidence during evaluation only in offer-sensitive mice. Our findings suggest that NAc core dopamine differentially reflects not only the decision made (accept vs skip), but also the confidence with which those decisions are made in more future-thinking offer-sensitive animals. That is, these data suggest that decision confidence regulates dopamine when these animals are evaluating decisions. After physiologically calibrating our optogenetic stimulation parameters uniquely for each animal, we also established a causal contribution of dopamine to decision confidence in offer-sensitive mice with more deliberative, future-thinking phenotypes.

In our stimulation study, we sought to augment dopamine dynamics in the offer zone in order to probe the extent to which this would alter evaluation of the offer. In doing so, we expected to see possible changes in the rate of accept and skip behaviors. To our surprise, stimulation did not influence the probability of accepting or skipping an offer, rather it altered the degree of confidence with which future-thinking animals made their decisions, and in the direction we would expect given our dLight recordings. This interpretation was further reinforced when animals reduced their rate of change-of-mind quitting in the wait zone following stimulation in the offer zone. Optogenetic stimulation yielded the greatest effects when stimulating at less preferred flavors, likely because behavior at less preferred flavors was more labile and amenable to alteration. These findings suggest that dopamine augmentation increased confidence in real-time during evaluation and these effects carried over into re-evaluation, where stimulated animals remained more committed to their decision.

Dopamine dynamics have been widely investigated in the context of decision-making^43,44^, with some evidence of individual differences related to stimulus-reward learning^14^. However, in these studies, decision-making is represented as a discrete event occurring at the moment at which an agent makes a choice. In contrast, most decision-making in the wild is a continuous and iterative process, encompassing evaluation before and re-evaluation after the choice. The evaluation of options often involves deliberation to accumulate confidence and arrive at decisions, while re-evaluation of decisions often involves counterfactual reasoning to imagine alternate outcomes.

Temporal difference reinforcement learning (TDRL) reward-prediction error (RPE) has been the prevailing theory regarding dopamine as a regulator of learning and decision-making^45–48^. While other theoretical proposals have been made that profoundly challenge the explanatory power of TDRL^41,49,50^, the behavioral procedures used to test these theories have been limited in their ability to capture important decision-making factors like confidence or counterfactual valuation. Our data suggest that dopamine causally regulates decision-making factors like confidence, counterfactual evaluation, and change-of-mind re-evaluation. Our work does not seek to overturn or contradict the classic TDRL model. Rather, we intend to make the case that the TDRL model should be expanded to include counterfactual prediction terms that capture change-of-mind processes during learning and decision-making. Just as an agent’s belief state can influence reward prediction computations^51,52^, so too, we argue, can an agent’s counterfactual reasoning. Thus, in addition to accounting for actual and expected outcomes, the model may benefit from integrating counterfactual predictions about what could have been. The formative TDRL algorithms calculate value dependent on the actual state of the world. This is learned by computing the difference between actual and predicted changes in value across each state. While this takes into account the agent’s expectations and actual values, there is no term in the standard model that considers alternative (counterfactual) values, as suggested by “economics of regret” models^20,53,54^. In cases where there are limited resources, competing options, or conflicting motivations, the value of each state may depend on both actual and counterfactual representations. Therefore, representing this in more modern conceptions of TDRL may account for discrepancies that original TDRL learning cannot explain.

Questions of change-of-mind and re-evaluation are usually discussed in terms of cognitive processes of regret^54^. Regret is distinct from disappointment — regret arises from mistakes of one’s own agency, while disappointment reflects unexpected losses^21,22,55^. RPE gives access to the latter through negative prediction errors, but dopamine’s contributions to the former remain unstudied. Our data suggest that dopamine plays a causal role in these questions of regret, and suggest future studies directed specifically at questions of regret may be particularly informative. More recent evidence highlighting the role of dopamine in signaling causal associations^50^, policies^41^, or perceived saliency^49^ provide compelling evidence to challenge RPE, but do not assess these complex decision-making strategies that take into account counterfactual outcomes such as regret and re-evaluation. Our findings suggest that dopamine regulates confidence in future choices and alters the re-evaluation of decisions that culminate in a change-of-mind. Furthermore, we found that the dopamine signals both reflected this information and causally modulated behavior in a manner related to the strategies individual animals used to achieve their goals.

We found that the effects of dopamine manipulation depended on the degree to which the animal considered the future. Overall, our work supports the theory that dopamine is not a monolith: the extent to which dopamine reflects cognitive processes like deliberation and re-evaluation depend on the strategies and means by which the agent makes decisions as well as the phase of the decision-making process (i.e., evaluation versus re-evaluation). Our findings regarding dopamine dips during quits also add to a literature that has largely focused on increases in dopamine levels, due to limitations in quantifying decreases in dopamine with classic methods like fast-scan cyclic voltammetry. When decreases have been reported, they have been associated with external noxious stimuli^56–58^ or externally-driven disappointment^13,59–61^, but never to internally-driven cognitive events such as re-evaluation and change-of-mind. Our data indicate that dips and rebounds in dopamine can encode unique and distinct aspects of counterfactuals and enable self-directed re-evaluation. Further, we found that dopamine manipulations interacted with confidence, which took into account both actual and counterfactual outcomes. By unveiling distinct decision-making strategies, we find that mesolimbic dopamine conveys information about past and future value and scales with decision confidence, specifically for behavioral strategies that depend on computations related to future outcomes. These individual differences in the fundamental operation of mesolimbic dopamine could present unique individual vulnerabilities to dopamine dysfunction and associated neuropsychiatric conditions^14,62–66^.

## METHODS

### Animals

All procedures were approved by the Institutional Animal Care and Use Committee at the University of Minnesota. Experiments were performed using comparable numbers of both female and male mice^28^. DAT-IRES-Cre transgenic mice^67^ were originally obtained from The Jackson Laboratory (JAX Stock #006660), and maintained on a C57BL/6J genetic background by breeding in-house. Following stereotaxic surgery at 8-12 weeks of age, mice were singly housed on a 14:10 light:dark cycle.

### Behavior

#### Pellet Training

Mice were introduced to flavored pellets 1 week prior to the start of their training. During this pre-training period, mice were transferred from regular rodent chow to a diet consisting of Bio-Serv full nutrition dustless precision pellets consisting of equal parts of chocolate, banana, grape, and plain flavored pellets (Bio-Serv product #F05301, #F0071, #F0079, #F07122). A free-feeding baseline weight was recorded as the average weight on three consecutive days of ad libitum pellet feed. Afterwards, daily feed was reduced by 0.5 g per day across four days. The day prior to beginning training, mice were introduced to the maze and given 15 minutes to explore the feeding sites. Each of the four feeders was filled with a specific flavor of pellets and surrounded by spatial cues to allow mice to become familiar with each restaurant.

#### Restaurant Row

Mice were run at the same time daily across consecutive days to maintain stable weights and motivational states. Training consisted of hour-long sessions of foraging on the maze in 4 stages. Stage 1 occurred on days 1-7 during which all offers were 1 second (associated tones = 4000 Hz, 500 msec). Stage 2 occurred on days 8-12 during which offers ranged from 1-5 seconds (associated tones ranged from 4000 – 5548 Hz, 500 msec). Stage 3 occurred on days 13-17 during which offers ranged from 1-15 seconds (associated tones ranged from 4000 – 9418 Hz, 500 msec). Stage 4 occurred on days 18+ during which offers ranged from 1-30 seconds (associated tones ranged from 4000 – 15223 Hz, 500 msec). Offers were pseudorandomly selected such that all offer lengths were sampled and then reshuffled independently for each flavor. Each offer tone was presented when mice entered into the offer zone in a counter-clockwise direction, and repeated each second until mice either accepted or skipped the offer. After accepting the offers, the countdowns in the wait zone decreased in pitch in 387 Hz steps once per second until the countdown ended at 4000 Hz, at which point a uniquely-flavored pellet was dispensed using a Med Associates dispenser. Any pellets that were not consumed were flushed using mini-servos to prevent mice from returning to uneaten rewards at leisure. Four Audiotek speakers were placed by each restaurant to provide local sound. Behavioral tracking and programming were conducted using a Logitech HD Webcam positioned above the maze and AnyMaze software. Pre and post weights were taken for each animal and small portions of post-training feed were given to maintain body weights at ∼80-85% of free feeding baseline. Photometry recordings and optogenetic manipulations were performed on the same maze as all training.

### Fiber Photometry

#### Surgery

Under ketamine:xylazine anesthesia (cocktail 100:10 mg/kg), holes were drilled above the NAc core (AP +1.35, ML +/- 2.13; DV -4.3, at a 10° angle from center) and VTA (AP -2.9, ML +/- 0.4; DV -4.55). Using a 33-gauge Hamilton syringe, 0.5 µL of AAV9-CAG-dLight1.3b (Addgene plasmid #125560; a gift from Lin Tian) and AAVdj-hSyn-FLEX-ChrimsonR (Addgene plasmid #62723; a gift from Edward Boyden) were injected at a rate of 0.1 µL/min bilaterally into the NAc core and VTA, respectively. After allowing 10 minutes for viral diffusion, the syringe was retracted slowly at a rate of 1 mm/min. Fiber-optic cannula (400 µm, Doric Lenses: MFC_400/430-0.48_6mm_MF1.25_FLT) were implanted 0.05 mm dorsal to the injection site at a 10° angle targeting the NAc core, and secured to the skull using jeweler’s screws and cured dental resin (Geristore). Virus was incubated for at least four weeks to allow for sufficient expression of dLight and opsin transport to mesolimbic dopamine terminals.

#### Data Collection

Dopamine dynamics were measured using a Tucker Davis Technologies RZ5P fiber photometry processer. Blue (470 nm) and violet (405 nm) LEDs (ThorLabs) were modulated at distinct carrier frequencies (531 Hz and 211 Hz, respectively) for dLight excitation and isosbestic control. LED output power was maintained between 50-75 μW. Signals were filtered through a fluorescence mini cube (Doric Lenses) and measured with a femtowatt photodetector (Newport), sampled at 6.1 kHz. The distal end of the cable was coupled to a fiber optic patch cord (400 µm, 0.48 NA, Doric Lenses) which connected to fiberoptic ferrules implanted in animals. Recordings hemispheres were counterbalanced across animals. Behavioral sessions were aligned to fiber photometry recordings using TTL signals sent from Anymaze to the RZ5P.

#### Processing

All recorded signals were analyzed offline. Changes in dLight fluorescence were measured by fitting the 405 nm isosbestic signal to the 470 nm signal and calculating dF/F ([470 nm signal-fitted 405 nm signal]/ [fitted 405 nm signal]). The output of this processing step effectively corrects for photobleaching as well as movement artifact, and was analyzed without any further rolling averages or smoothing. For behavioral event related analyses, signals were aligned to relevant behavioral timepoints and averaged within subjects across trials, then between subjects across days.

### Optogenetic validations

We used a five-port minicube (Doric Lenses) to filter excitation and emission channels for combined fiber photometry and optogenetic manipulations. For stimulation using ChrimsonR, a Master-8 (AMPI) was used to drive a 589 nm laser (Opto Engine LLC) to generate 0, 5, 10, 15, and 20 pulses (5 ms) at 20 Hz to stimulate VTA terminals in NAc expressing ChrimsonR, while simultaneously recording dLight1.3b responses through the RZ5P as described above. Five technical replicates were conducted for each parameter and an interval of 30 seconds was maintained between each stimulation. Laser power was set at 3-4 mW output to produce dopamine responses with amplitudes similar to the largest endogenous response observed after earning reward. Unique stimulation parameters were selected for each individual animal to ensure that dopamine responses were standardized across mice, and resembled physiologic levels of dopamine seen during task performance.

To confirm that halorhodopsin effectively reduced dopamine release in the NAc (Fig. 6b-d), we delivered light (589 nm, 2 seconds continuous, 6-8 mW) coincident with food pellet delivery, when a robust increase in dopamine signal is normally observed during reward receipt. We again used a five-port minicube (Doric Lenses) to filter excitation and emission channels for combined fiber photometry and optogenetic inhibition. Using closed-loop behavioral tracking through Anymaze, we delivered light on half of trials when animals earned a pellet across two training days.

### Optogenetic manipulations during behavior

After completing training, mice underwent a series of optogenetic stimulation testing days. An Anymaze Optogenetic Interface was coupled to a 589 nm laser (Opto Engine LLC) to interface between the behavioral software and light source. Closed-loop behavioral tracking through Anymaze allowed for laser activation at precise behavioral timepoints.

For ChrimsonR experiments, light (individually-tailored number of pulses, 20 Hz, 5 ms) was delivered immediately upon crossing into the offer zone and terminated if the animal accepted or skipped the trial. Mice received stimulation within the offer zone on 50% of offers above 15 seconds, selected randomly across 1) all flavors, 2) less preferred flavors, and 3) more preferred flavors on different days. The order of these testing conditions was counterbalanced between animals. Stimulation days were interleaved with non-stimulation behavioral days. To control for the possibility that light alone was driving our observed effects, we mirrored light delivery conditions in two control experiments. First, we tested mice lacking Cre recombinase using the average stimulation settings for mice expressing Cre recombinase (11 pulses of light, 589 nm, 20 Hz, 3-4 mW), and delivered light in the offer zone on half of all trials at less preferred flavors. Second, we evaluated the effects of delivering this pulsed light pattern in a mismatched fashion to mice expressing halorhodopsin rather than ChrimsonR. We measured dLight1.3b signals during this mismatched pattern of stimulation, to confirm that pulsed light was not sufficient to engage the inhibitory effects of halorhodopsin (Extended Data Fig. 7e-g). We then delivered this mismatched stimulation during behavior, mirroring the conditions described above for mice expressing ChrimsonR, with light delivery in the offer zone on half of trials at less preferred flavors.

For halorhodopsin experiments, wait zone entry triggered continuous light delivery for up to 4 seconds during the countdown, to match the time course of the dopamine dips observed during quitting. If animals quit during the 4 seconds of light delivery, the laser would turn off. Mice received inhibition in the wait zone on 50% of offers above 15 seconds selected randomly across 1) all flavors, 2) less preferred flavors, and 3) more preferred flavors on different days. The order of these testing conditions was counterbalanced between animals. Inhibition days were interleaved with non-stimulation behavioral days. In condition (2), mice accepted offers at less preferred flavors with a low probability, so the number of trials with light delivery were too few for reliable analysis. We therefore report results only for light delivery at all flavors or more preferred flavors.

### Analysis

*Figure 1*: Laps and number of pellets earned were calculated for each day. Laps were defined as a full counterclockwise rotation around the maze with an offer at each restaurant. Number of pellets were separated by flavor rank each day. Time spent before quitting was calculated for each trial that resulted in a quit and averaged by rank. Proportion of trials resulting in a quit versus skip were calculated for each day. For change of mind analyses, we sought to behaviorally distinguish quitting from skipping. We conducted an analysis probing behavioral outcomes on the subsequent trial (next restaurant: “R+1”), based on whether the animal exhibited an immediate skip or a change-of-mind quit on the previous trial (“R”). Probability of accepting, reaction time, IdPhi, and lingering time were calculated at R+1 after skipping or quitting at R (Extended Data Fig. 1).

*Figure 2*: Offer sensitivity was quantified as the difference in probability of accepting a good offer (0-3 seconds) and probability of accepting a bad offer (27-30 seconds). After calculating this measure of offer sensitivity for each individual animal, we fit Gaussian mixture models with varying numbers of components to the distribution of offer sensitivity. The Akaike information criterion (AIC) was minimized by a model with two components providing quantitative evidence for a distinction between two decision-making phenotypes. We used this two-component Gaussian mixture model to separate “offer-sensitive” mice from “offer-insensitive” mice that exhibited little or no offer sensitivity.

Probability of accepting, skipping, quitting, and earning were calculated as a proportion of all offers (trials), separated by flavor preference rank. Differences in skipping offers among behavioral phenotypes was quantified as the probability of skipping a 1s offer subtracted from the probability of skipping for a 30s and separated by rank. Thresholds were calculated each day, for each animal and each flavor. For offer zone thresholds, we fit a sigmoid to the binary offer zone outcome (accept or skip) as a function of offer and calculated the point of inflection (i.e., the offer at which a particular animal would shift from accepting offers to skipping offers for a specific flavor). Using a similar approach, we calculated wait zone thresholds, fit to the wait zone outcomes (earn or quit), and calculated the offer at which a particular animal would shift from earning to quitting for a specific flavor.

*Figure 3*: Dopamine responses to offer were binned for short (1-5s), medium (6-15s), and long (16-30s) offers, and aligned to offer onset during mid-training (Days 30-37). Using a masking function, the data were restricted from the transition from the previous restaurant to the current trial’s offer zone to constrain our analysis to offer-related dopamine transients. Responses post offer were then averaged (0 to 3s) and normalized to a pre-offer period (-2 to -1s). Dopamine was then aligned to offer zone outcomes (accept and skip) and wait zone outcomes (earn and quit). Mean dopamine signals were calculated for accepts, skips, and quits by averaging dF/F during the pre-event period (-1 to 0s) or post-event period (0 to 2s) for earns, and normalizing to a preceding baseline (-2 to -1s).

*Figure 4*: Analysis of past value was conducted by calculating the value of the offer received earlier in the offer zone at the time of quit (Past Value = Threshold - Offer). Thresholds were calculated for each animal each day using the methods described above. Peri-event (quit and skip) dopamine signals were grouped in three bins: low past value (</= -14), medium past value (</= 0), and high past value (> 0). Past time spent was analyzed by calculating the time the animal invested before quitting and was grouped in three bins: low time spent (</= 4s), medium time spent (4s to </=6s), and high time spent (>6s). Mean changes in dopamine to value remaining and time remaining were calculated by averaging the dopamine signal during the dips (0.5 to 1.5s) and normalizing to a pre-dip baseline (-3 to -2s).

*Figure 5*: Analysis of future value was conducted by calculating value remaining at the time of quit: Value Remaining (future value) = Threshold – Time Remaining in Countdown. Thresholds were calculated for each animal each day using the methods described above. Time remaining at countdown was defined as the number of seconds remaining in the countdown the animal would have had to wait to have earned reward and was calculated as Offer – Time to Quit. Peri-event dopamine signals were grouped in two bins: favorable future values (Value Remaining <0) and unfavorable future values (Value Remaining >0) and aligned to quits. Trials were randomly subsampled to match favorable and unfavorable quit types within group and offer-sensitive and offer-insensitive quits between groups. Future time remaining was analyzed by splitting data into low (</=15s) and high (>15s) time remaining. Mean changes in dopamine to value remaining and time remaining were calculated by averaging the dopamine signal during the dips (-1.5 to 0s) and normalizing to a pre-dip baseline (-3 to -2s).

*Figure 6*: Probability of quitting was calculated for each pass in the wait zone and average probability of quitting on offers above 15 seconds were compared on trials with and without light delivery. Difference scores were calculated between these trial types.

*Figure 7*: Absolute integrated angular velocity was calculated for each pass in the offer zone and used to signify IdPhi for each trial. Trials were split based on the z-score of their IdPhi values: zIdPhi > 0 were classified as VTE and zIdPhi < 0 were classified as non-VTE. IdPhi relates to path curvature, so we utilized a path analysis previously reported in studies characterizing tortuosity of blood vessels in the retina^68^. Dopamine signals were aligned to the point of maximum curvature and separated based on trial outcome (accept or skip). Mean dopamine was calculated as the average dF/F post-max curvature (0.5 to 1.5s) and normalized to a preceding baseline (-4 to -1s).

*Figure 8*: zIdPhi was calculated for each pass in the offer zone and average zIdPhi on offer above 15 seconds were compared on stimulated and matched non-stimulated trials. Difference scores were calculated between stimulated and non-stimulated cases. Probability of quitting was calculated for each trial and probabilities were compared on trials with and without light delivery in the offer zone. Difference scores were calculated between these trial types.

### Immunohistochemistry

After behavioral testing, mice were deeply anesthetized using Beuthanasia (200 mg/kg, intraperitoneal) and transcardially perfused with ice-cold PBS and 4% paraformaldehyde in PBS for 15 minutes. Brains were extracted and post-fixed overnight in 4% paraformaldehyde in PBS at 4° C. Coronal sections were collected at 50 µm thickness using a vibrating microtome (Leica Microsystems) for staining with immunohistochemistry. Nonspecific binding was blocked with 2% normal horse serum + 0.05% Tween-10 + 0.2% Triton-X100 in PBS overnight at 4° C. After washing, slices were then incubated in mouse anti-GFP (Invitrogen, A-11120, diluted 1:500) rabbit anti-RFP (Rockland, 600-401-379, diluted 1:1000) primary antibodies diluted in the same blocking solution for 24 hours at 4° C. After four rinses in PBS containing 0.1% Tween-20, sections were transferred to secondary antibody goat anti-mouse IgG Alexa 488 (Abcam, ab150115, diluted 1:1000) and donkey and rabbit Alexa 647 (Abcam, ab150075, diluted 1:1000) diluted in blocking solution for 24 hours at 4° C. Sections were rinsed three times using PBS with 0.1% Tween-20 then mounted on slides and coverslipped using DAPI mounting (ProLong Gold antifade reagent with DAPI, Lot #250196) and imaged using fluorescence microscopy (Leica, BZ-X Series). Fiber optic tip locations were mapped using the Paxinos and Franklin’s Mouse Brain Atlas^69^.

## Data Availability

Data will be posted to OSF after reviews are completed and a public version of the paper has been made available.

## Code Availability

Code will be posted to OSF after reviews are completed and a public version of the paper has been made available.

## Acknowledgments

This work was supported by grants from the National Institutes of Health (F30MH124404 to AK; P50MH119569 to A.D.R. and P.E.R.; R01DA048946 to P.E.R.); the J. B. Johnston Land Grant Chair in Neuroscience (A.D.R.); and the University of Minnesota’s MnDRIVE (Minnesota’s Discovery, Research, and Innovation Economy) initiative (P.E.R.). Some viral vectors used in this study were generated by the University of Minnesota Viral Vector and Cloning Core and the Center for Neural Circuits in Addiction (P30 DA048742). The University of Minnesota MnDRIVE Optogenetics Core provided technical support for fiber photometry experiments. We thank the University of Minnesota Mouse Behavior Core for use of their facilities to conduct behavioral tests. We also thank Dr. Brian Sweis and Dr. Mark Thomas for their insights and stimulating discussions, as well as Jacqueline Rouche and Hailie Bogenrief for enthusiastic technical assistance.

## Author contributions

Conceptualization: A.K., A.D.R., and P.E.R. Methodology: A.K., A.D.R., and P.E.R. Investigation: A.K. Analysis and visualization: A.K., A.D.R., and P.E.R. Funding acquisition and project administration: A.K., A.D.R. and P.E.R. Writing: A.K., A.D.R., and P.E.R.

## EXTENDED DATA FIGURES & LEGENDS

**Extended Data Fig. 1.**
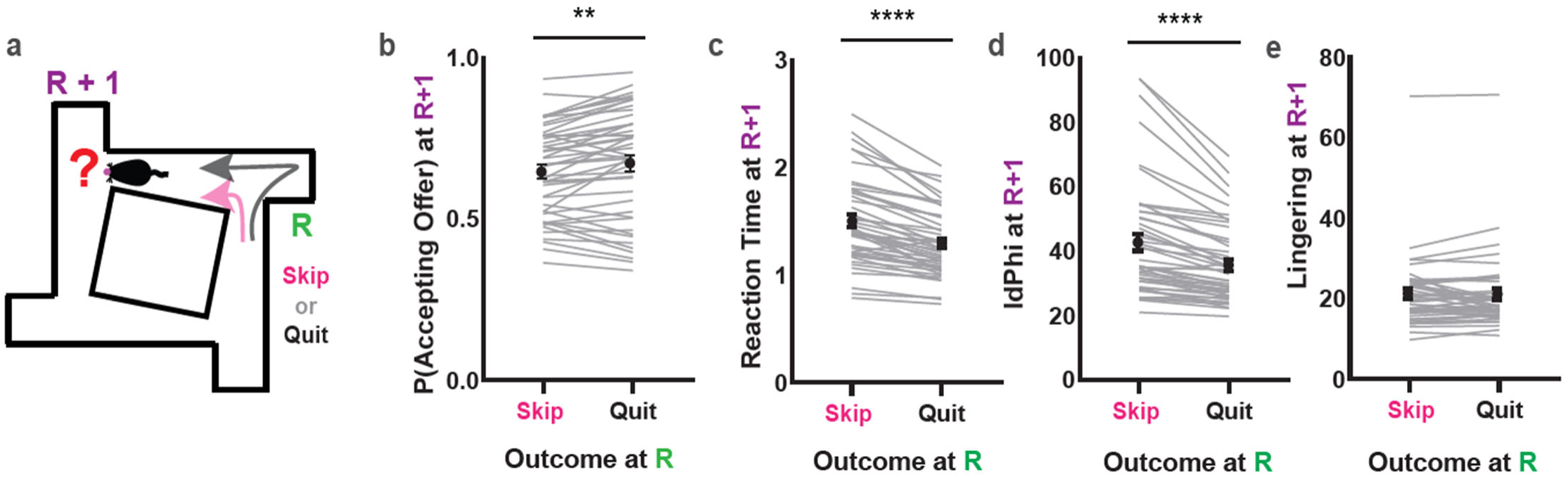
Behavioral differences between change-of-mind quitting versus skipping. **(a)**Analysis of behavior at the subsequent restaurant (R+1) after an animal skips or quits at the previous restaurant (R). **(b-e)** Probability of accepting offer (b), reaction time (c), IdPhi (d), and lingering time (e) at R+1 after skipping versus quitting at R. Data are mean +/- SEM for all panels (n=46). **p<0.01, ****p<0.0001, ANOVA main effect of Outcome; complete statistics are provided in Supplementary Tables.

**Extended Data Fig. 2.**
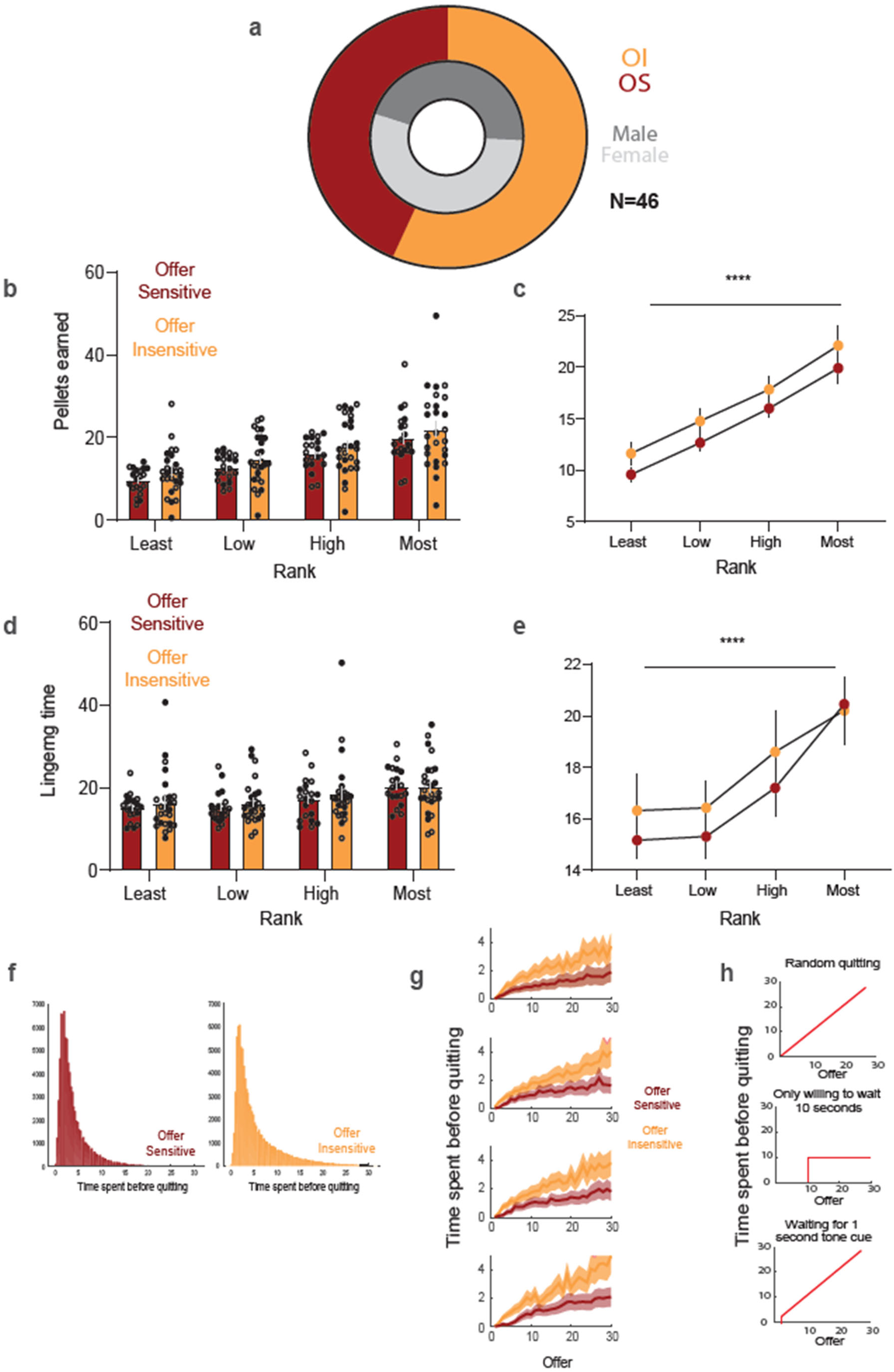
Characteristics of offer-sensitive and offer-insensitive mice. **(a)** Proportions of male and female offer-sensitive and offer-insensitive mice. **(b)** Pellet earnings for individual mice by rank. **(c)** Average pellet earnings by rank. **(d)** Lingering time for individual mice by rank, **(e)** Average lingering time by rank. **(f)** Distribution of time spent before quitting in offer-sensitive and offer-insensitive mice. **(g)** Time spent before quitting as a function of offer, separated by flavor rank. **(h)** Illustration of alternate strategies that are not being employed by mice of either phenotype. Neither phenotype was quitting randomly (*top*), only willing to wait a fixed amount of time (e.g., 10 seconds; *middle*); or waiting for a specific tone to cue them to quit (e.g., 1 second; *bottom*). ****p<0.0001, ANOVA main effect of Rank; complete statistics are provided in Supplementary Tables.

**Extended Data Fig. 3.**
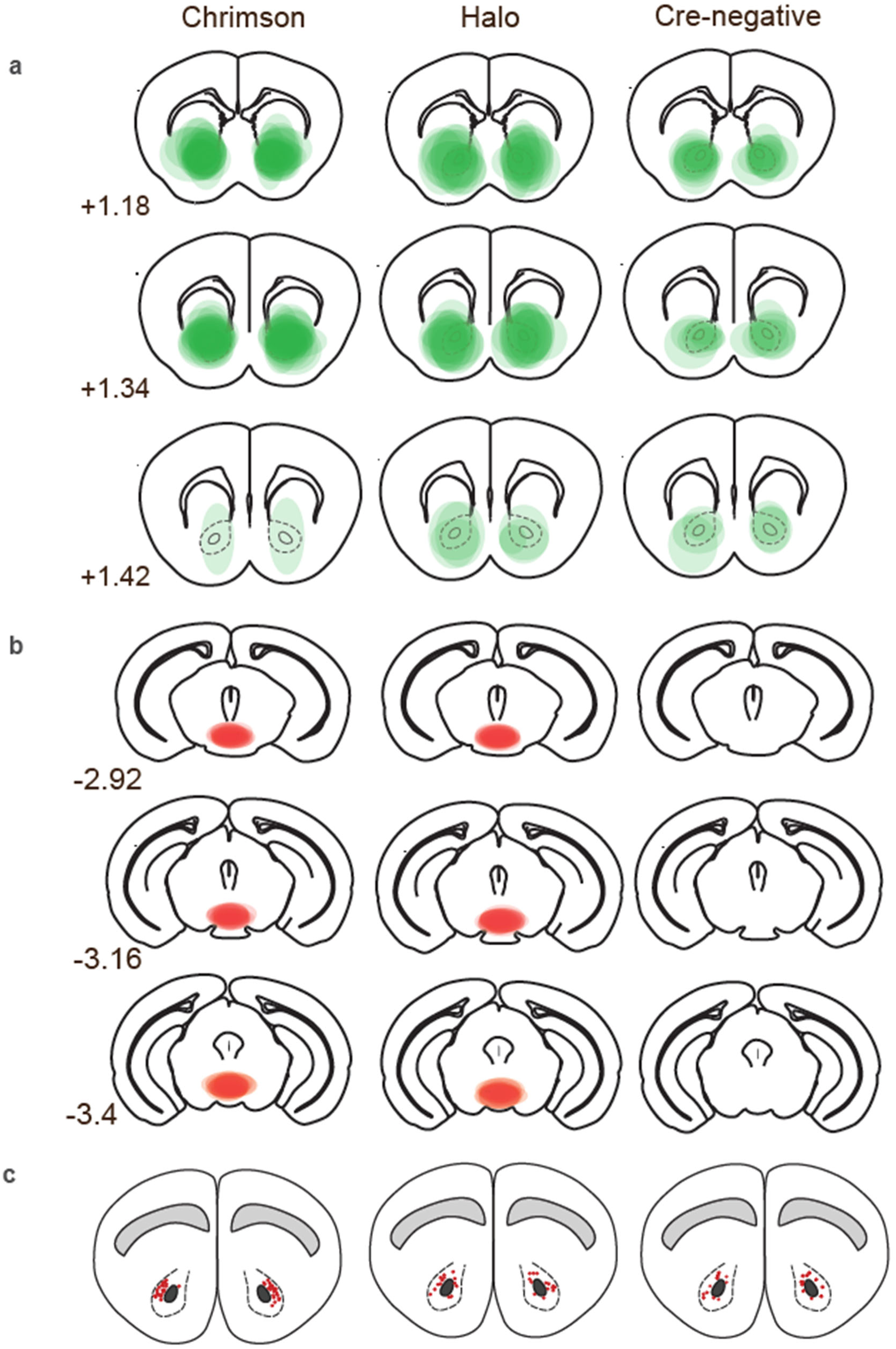
Anatomical maps of virus expression and optic fiber placements. **(a)** dLight1.3b viral expression in the nucleus accumbens. **(b)** ChrimsonR or eNpHR3.0 expression in VTA. **(c)** Fiber optic placement in nucleus accumbens core. Sequential columns show groups injected with ChrimsonR in the VTA, halorhodopsin in VTA, and the Cre-negative control group.

**Extended Data Fig. 4.**
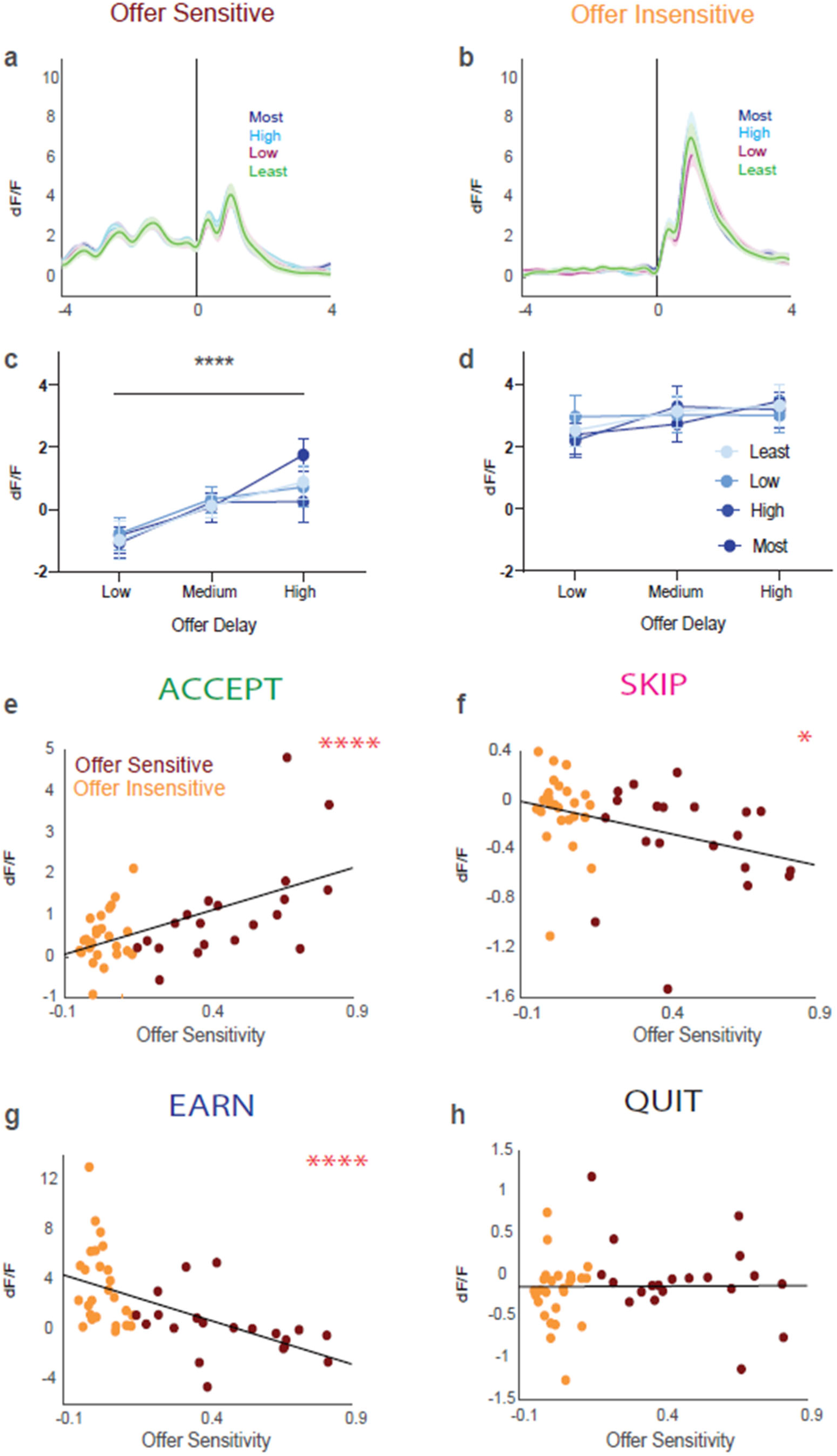
Dopamine responses to earning pellets of different flavors and correlations between offer sensitivity and dopamine dynamics during main trial events. **(a to b)** Dopamine signal aligned to pellet delivery in offer-sensitive (a) and offer-insensitive (b), separated by individual preference: most, high, low, and least (descending order). **(c to d)** Dopamine responses to earning as a function of both flavor preference and offer delay. Data are mean +/- SEM. **(e-h)** Average dopamine signal for accept (e), skip (f), earn (g), and quit (h) as a function of offer sensitivity. Yellow circles represent offer-insensitive mice (n=26), while red circles represent offer-sensitive mice (n=20). *p<0.05, ****p<0.0001, ANOVA main effect of Offer Delay (c) or Pearson correlation coefficient (e to g); complete statistics are provided in Supplementary Tables.

**Extended Data Fig. 5.**
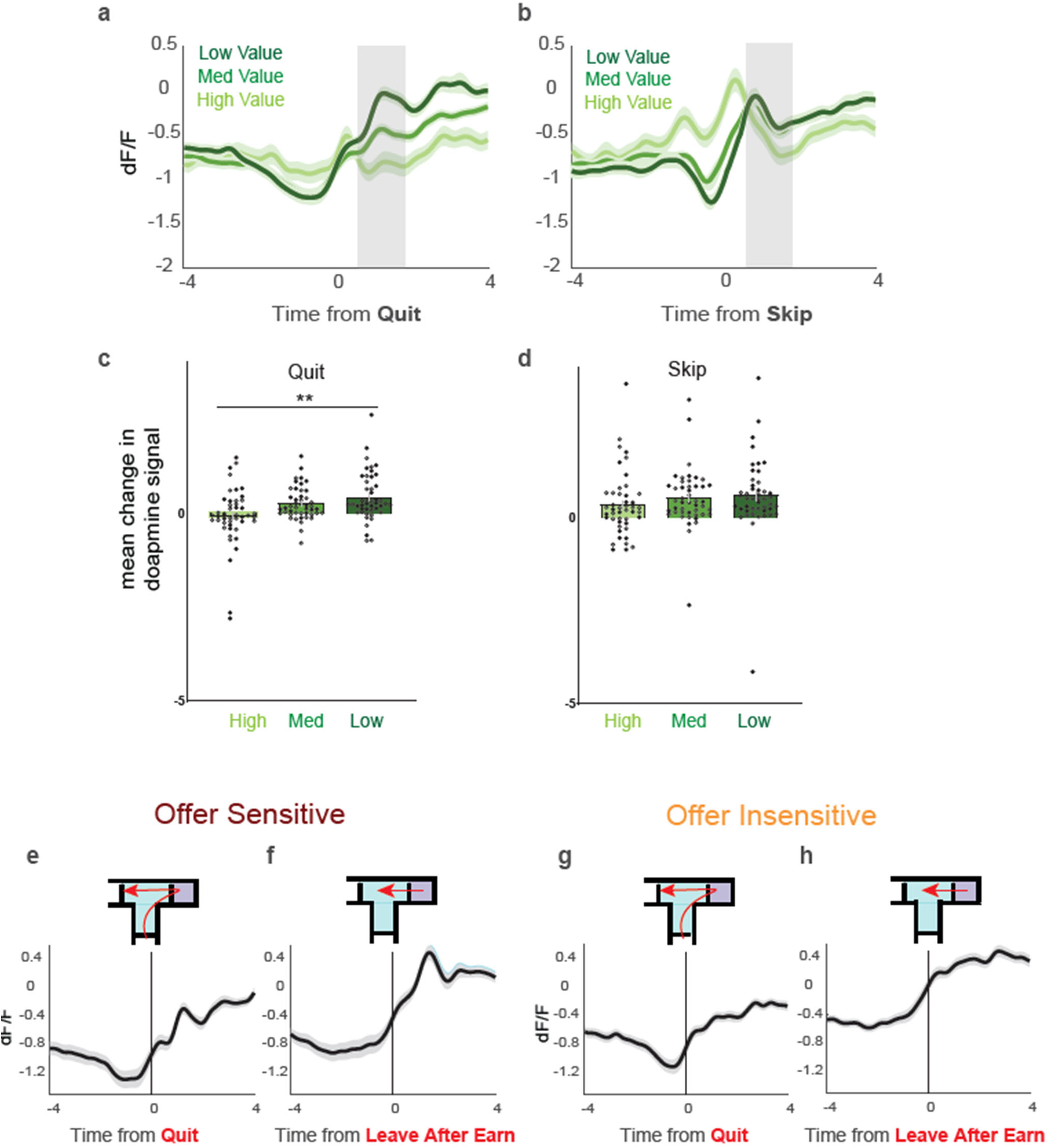
Dopamine dips and rebounds are specific to change-of-mind quitting. (**a and c**) Dopamine dynamics after the decision (time indicated by gray bar) scale inversely with past value following (but not preceding) a change-of-mind quit decision. (**b and d**) Dopamine dynamics scale inversely with past value preceding (but not following, time indicated by gray bar) a skip decision. **(e-h)** Dopamine signals differed between motorically identical but psychologically distinct acts of leaving the wait zone after quitting (e and g) versus earning (f and h), in both offer-sensitive (e-f) and offer-insensitive (g-h). Data are mean +/- SEM for all panels (n=46); open and filled circles represent female and male mice, respectively. **p<0.01, ANOVA main effect of Value; complete statistics are provided in Supplementary Tables.

**Extended Data Fig. 6.**
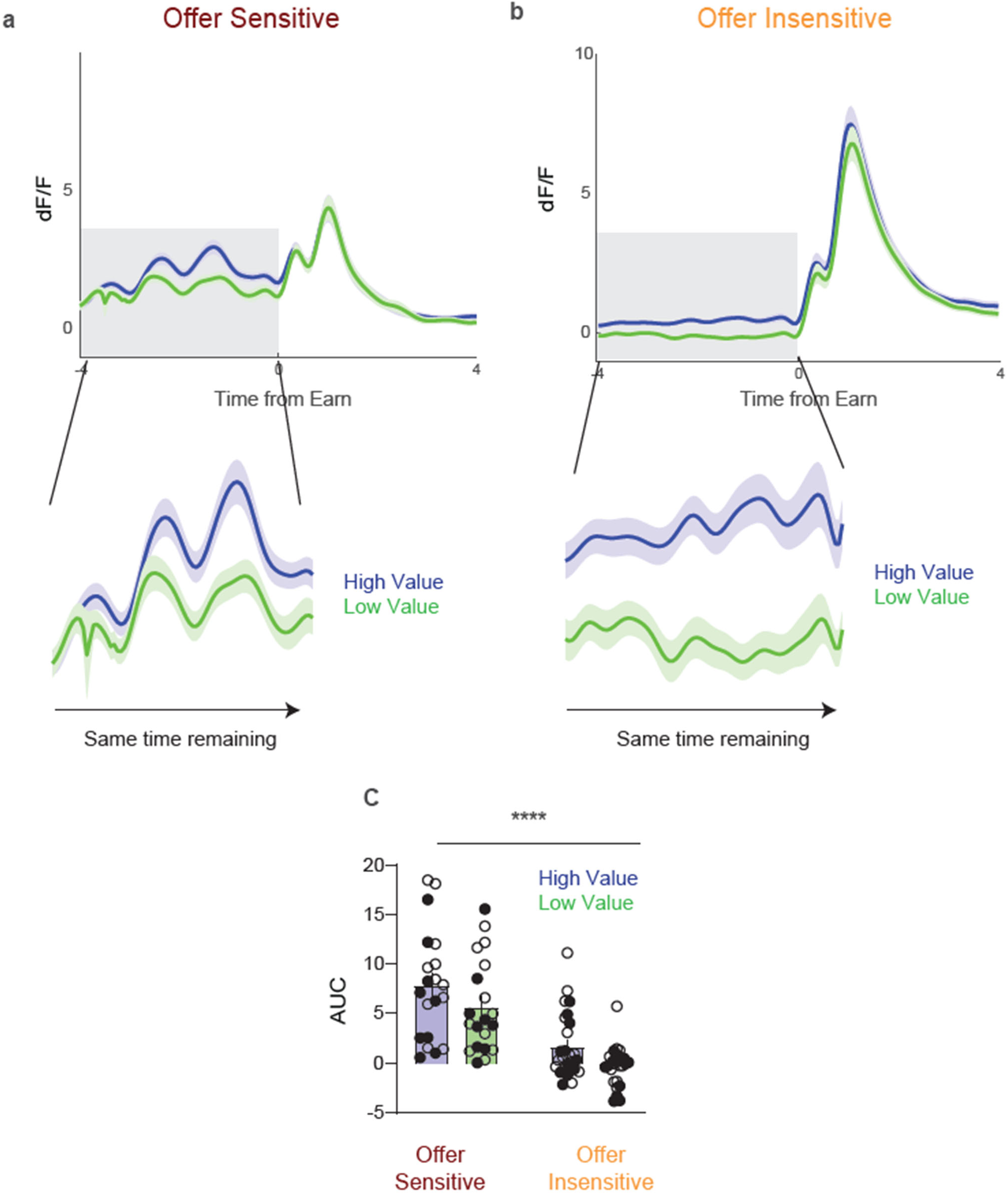
Dopamine reflects value and not time to earn. (**a**) Offer-sensitive and (**b**) offer-insensitive mice display elevated dopamine during the countdown on higher value offers, despite identical time to reward. (**c**) Area under the curve for high- and low-value offers over equal windows of time to reward (4 seconds). Data are mean +/- SEM for all panels; open and filled circles represent female and male mice, respectively. ****p<0.01, ANOVA main effect of value; complete statistics are provided in Supplementary Tables.

**Extended Data Fig. 7.**
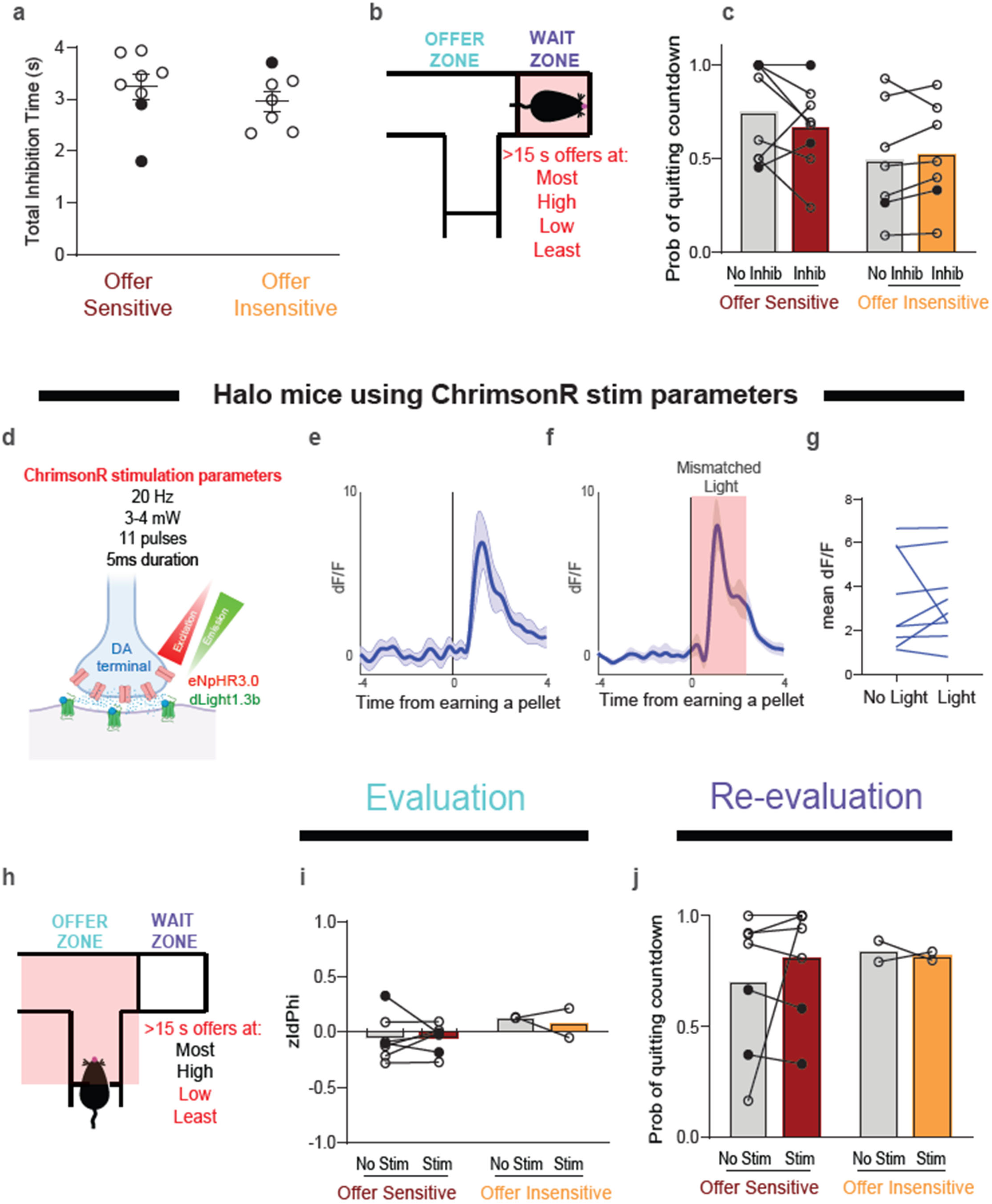
Control conditions for mice expressing halorhodopsin. **(a)** Average total duration of inhibition in wait zone for each trial (n=8/7 for offer-sensitive/insensitive). (**b**) Schematic showing wait zone locations (red shading and text) where light was delivered on half of offers > 15 s. (**c**) Probability of quitting on trials without light delivery (No Inhib) and with light delivery (Inhib) at all flavors. **(d)** Illustration of “mismatched” light delivery paremeters (589 nm, 20 Hz, 3-4 mW) for mice expressing halorhodopsin in mesolimbic dopamine terminals. **(e-g)** Dopamine response to earning a food pellet in the absence **(e)** and presence **(f)** of mismatched light delivery, along with the change in peak dopamine signal under each condition **(g)**. **(h)** Schematic showing mismatched light delivery in the offer zone (red shading and text). **(i)** zIPhi in the offer zone on stimulation trials with light delivery (“Light”) and control trials with no light delivery (“No Light”). **(g)** Probability of quitting in the wait zone with and without light delivery. Data are mean +/- SEM for all panels (n=7/2 for offer-sensitive/insensitive); open and filled circles represent female and male mice, respectively. Complete statistics are provided in Supplementary Tables.

**Extended Data Fig. 8.**
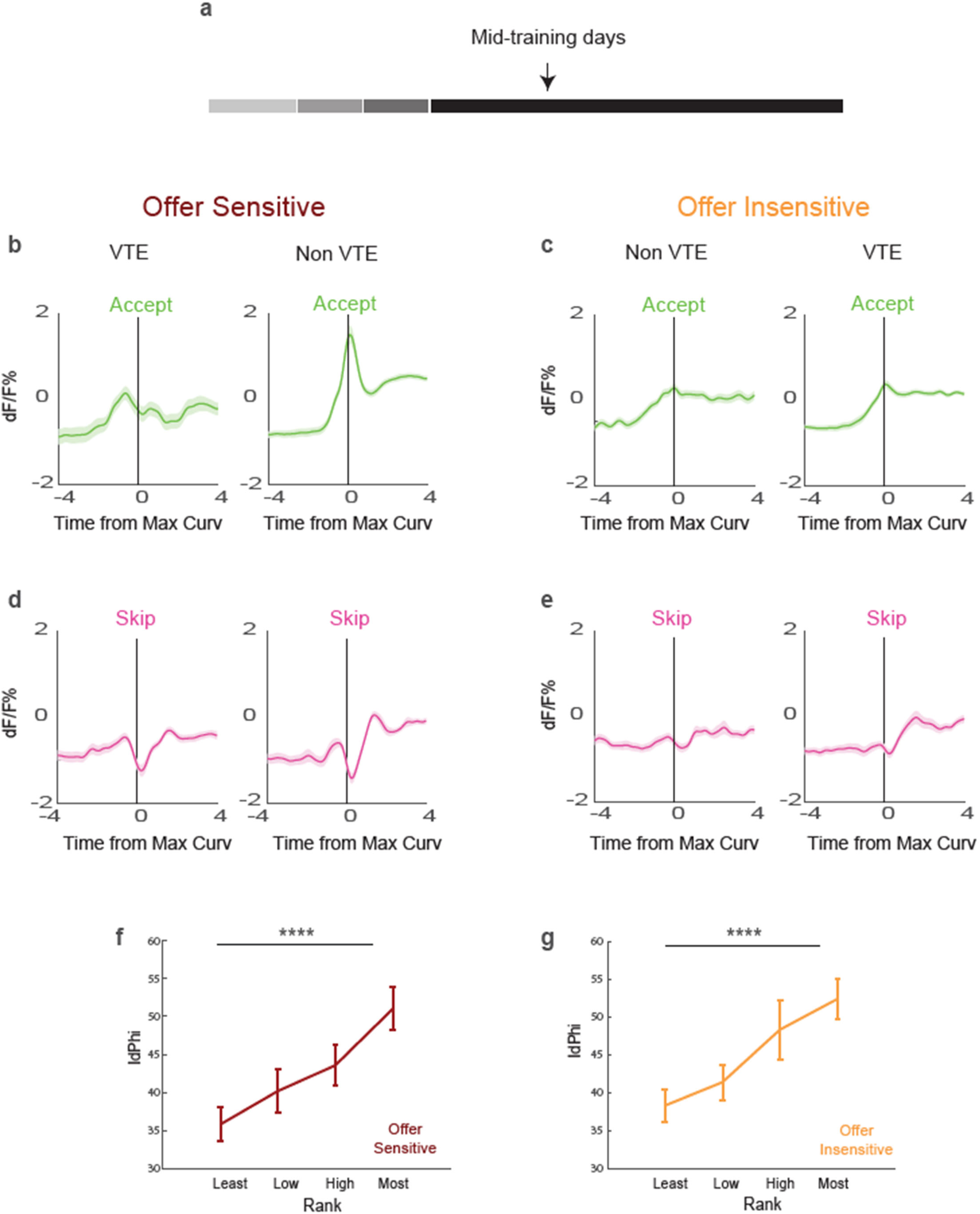
Dopamine dynamics relate to decision confidence earlier in training, follow similar trends as late training, and scale with flavor preference. **(a)** Mid training recording days. (**b)** Offer-sensitive and **(c)** offer-insensitive dopamine dynamics during accepts aligned to time at which maximum path curvature occurs on Non-VTE (right) and VTE (left) trials; **(d)** Offer-sensitive and **(e)** offer-insensitive dopamine dynamics during skips aligned to time at which maximum path curvature occurs on Non-VTE (right) and VTE (left) trials. **(f-g)** IdPhi scaled with flavor preference in both offer-sensitive (f) and offer-insensitive mice (g), such that there was greater VTE at more preferred flavors. Data are mean +/- SEM for all panels. ****p<0.0001, ANOVA main effect of Rank; complete statistics are provided in Supplementary Tables.

**Extended Data Fig. 9.**
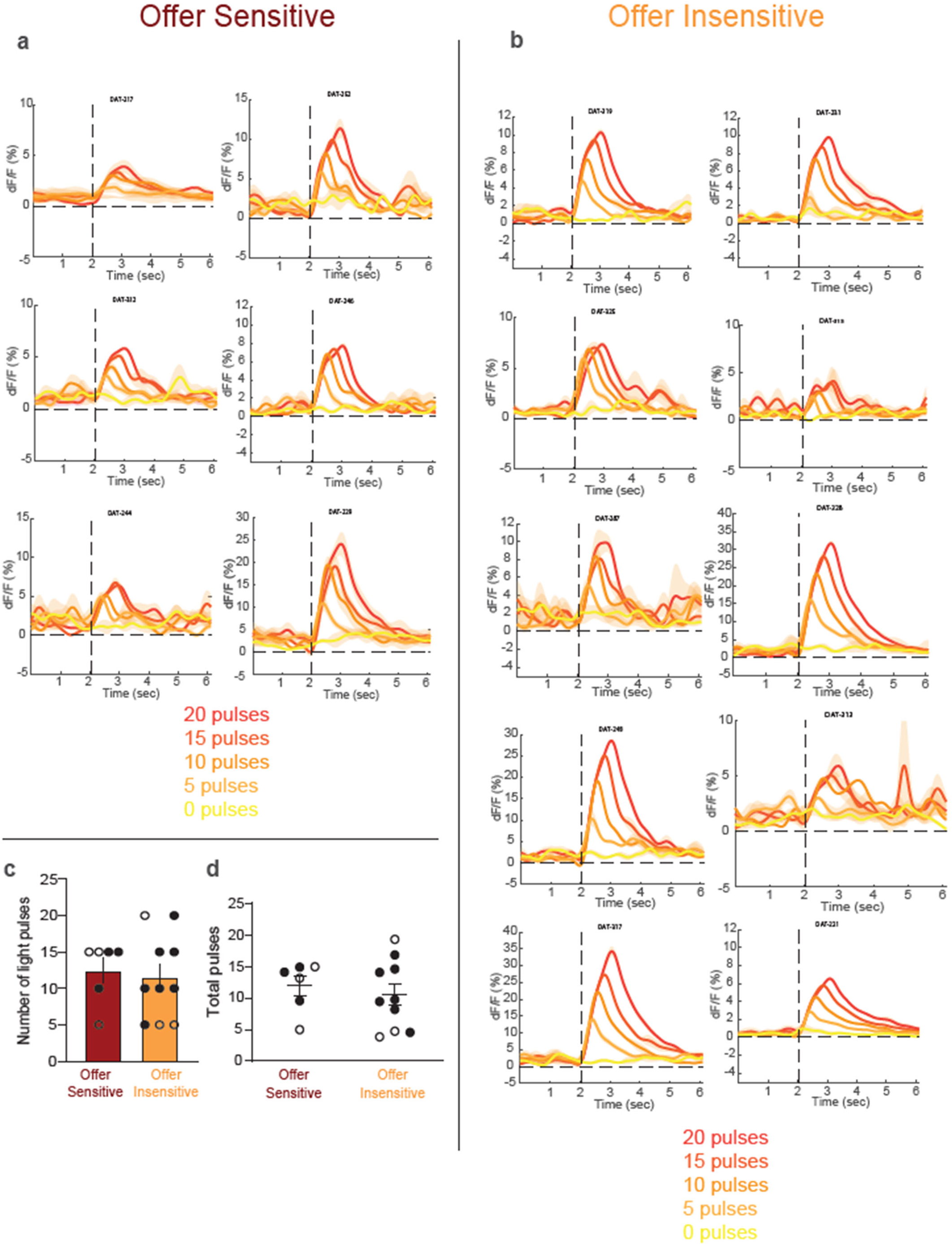
Individually-tailored optogenetic stimulation responses. **(a to b)** Optogenetic stimulation-response curves for each offer-sensitive (a) and offer-insensitive (b) mouse using the following parameters: pulse number 0, 5, 10, 15, or 20; wavelength 589 nm; frequency 20 Hz. **(c)** Stimulation parameters did not differ on average between offer-sensitive and offer-insensitive mice. **(d)** Average total number of pulses delivered in the offer zone for each trial did not differ (n=6/10 for offer-sensitive/insensitive). Data are mean +/- SEM for all panels; open and filled circles represent female and male mice, respectively. Complete statistics are provided in Supplementary Tables.

**Extended Data Fig. 10.**
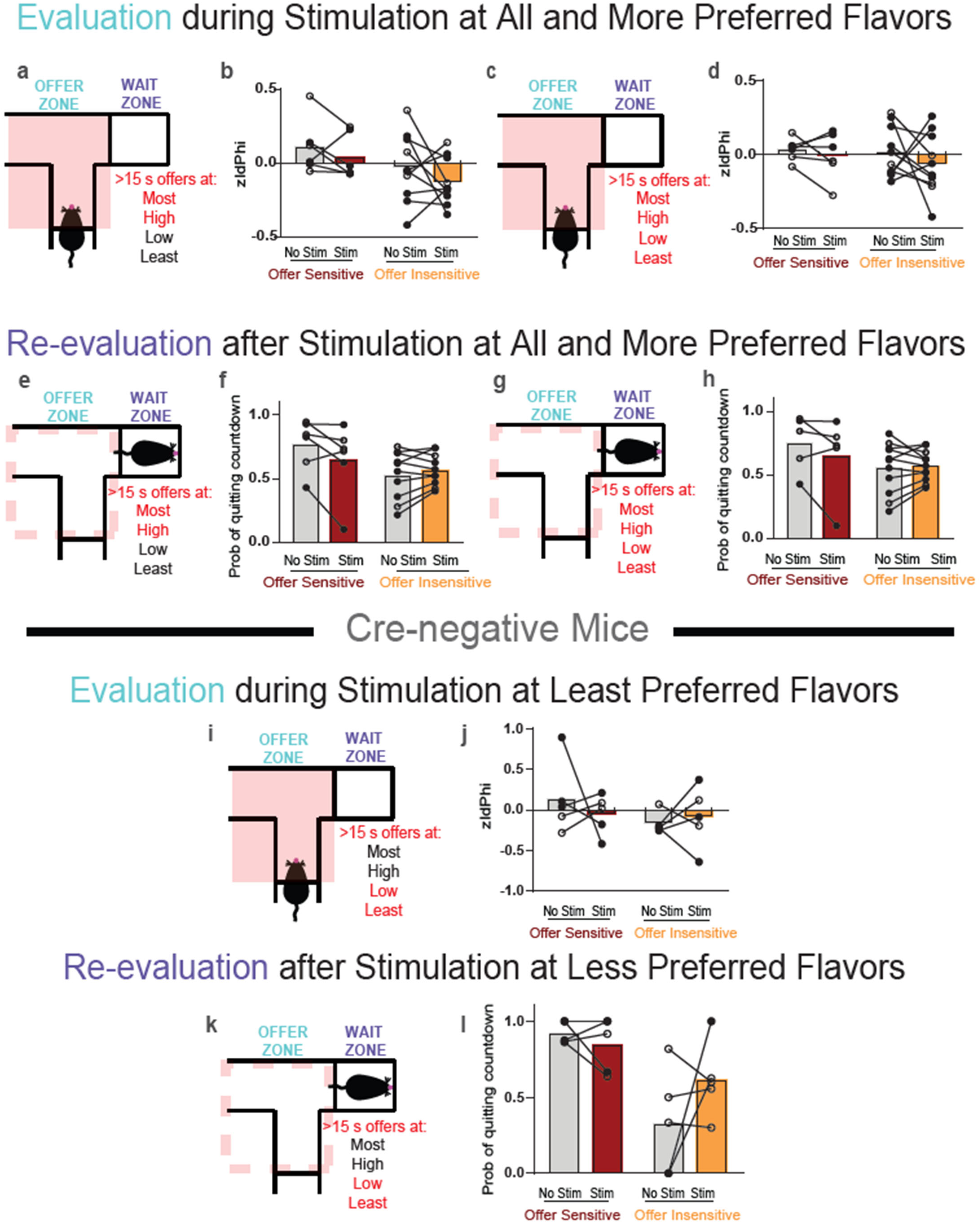
Specificity in the effects of optogenetic enhancement of dopamine release. **(a-d)** Schematics showing offer zone locations (red shading and text) where light was delivered on half of all offers > 15 s (a and c), with corresponding zIdPhi on trials without (No Stim) and with light delivery (Stim) at indicated flavors (b and d). **(e-h)** Schematic showing behavior in the wait zone after light delivery in the offer zone (dotted red line) had ended (e and g), with corresponding probability of quitting on trials after no light delivery (No Stim) or light delivery (Stim) at indicated flavors (f and h). **(i)** Schematic showing offer zone locations (red shading and text) where light was delivered to Cre-negative control mice on half of all offers > 15 s. **(j)** zIPhi on trials without (No Stim) and with light delivery (Stim) at less preferred flavors. **(k)** Schematic showing behavior of Cre-negative control mice in the wait zone after light delivery in the offer zone (dotted red line) had ended. **(l)** Probability of quitting on trials after no light delivery (No Stim) or light delivery (Stim) at less preferred flavors. Open and filled circles represent female and male mice, respectively. Complete statistics are provided in Supplementary Tables.

## MAIN REFERENCES

1. Van Hoeck, N., Watson, P. D. & Barbey, A. K. Cognitive neuroscience of human counterfactual reasoning. Front Hum Neurosci (2015) doi:10.3389/fnhum.2015.00420.

2. Kishida, K. T., et al. Subsecond dopamine fluctuations in human striatum encode superposed error signals about actual and counterfactual reward. Proc Natl Acad Sci U S A 113, 200–205 (2016).

3. Boldt, A., Schiffer, A.-M., Waszak, F. & Yeung, N. Confidence Predictions Affect Performance Confidence and Neural Preparation in Perceptual Decision Making. doi:10.1038/s41598-019-40681-9.

4. Zylberberg, A., Wolpert, D. M. & Shadlen, M. N. Counterfactual Reasoning Underlies the Learning of Priors in Decision Making. Neuron 99, 1083–1097.e6 (2018).

5. Roese, N. J. & Olson, J. M. Self-Esteem and Counterfactual Thinking. J Pers Soc Psychol (1993) doi:10.1037/0022-3514.65.1.199.

6. Sanna, L. J., Meier, S. & Turley-Ames, K. J. Mood, self-esteem, and counterfactuals: Externally attributed moods limit self-enhancement strategies. Soc Cogn 16, 267–286 (1998).

7. Josephs, R. A., Larrick, R. P., Steele, C. M. & Nisbett, R. E. Protecting the Self From the Negative Consequences of Risky Decisions. J Pers Soc Psychol (1992) doi:10.1037/0022-3514.62.1.26.

8. Mohebi, A., et al. Dissociable dopamine dynamics for learning and motivation. Nature 570, 65–70 (2019).

9. Sharpe, M. J., et al. Dopamine transients are sufficient and necessary for acquisition of model-based associations. Nat Neurosci 20, 735–742 (2017).

10. Steinberg, E. E., et al. A causal link between prediction errors, dopamine neurons and learning. Nat Neurosci 16, 966–973 (2013).

11. Hamid, A. A., et al. Mesolimbic dopamine signals the value of work. Nat Neurosci 19, 117–126 (2015).

12. Howe, M. W., Tierney, P. L., Sandberg, S. G., Phillips, P. E. M. & Graybiel, A. M. Prolonged dopamine signalling in striatum signals proximity and value of distant rewards. Nature 500, 575–579 (2013).

13. Schultz, W., Dayan, P. & Montague, P. R. A neural substrate of prediction and reward. Science (1979) 275, 1593–1599 (1997).

14. Flagel, S. B., et al. A selective role for dopamine in stimulus-reward learning. Nature 469, 53–59 (2011).

15. Wassum, K. M., Ostlund, S. B., Loewinger, G. C. & Maidment, N. T. Phasic mesolimbic dopamine release tracks reward seeking during expression of pavlovian-to-instrumental transfer. Biol Psychiatry (2013) doi:10.1016/j.biopsych.2012.12.005.

16. Aitken, T. J., Greenfield, V. Y. & Wassum, K. M. Nucleus accumbens core dopamine signaling tracks the need-based motivational value of food-paired cues. J Neurochem 136, 1026–1036 (2016).

17. Saunders, B. T., Richard, J. M., Margolis, E. B. & Janak, P. H. Dopamine neurons create Pavlovian conditioned stimuli with circuit-defined motivational properties. Nat Neurosci 21, 1072–1083 (2018).

18. Taira, M., et al. Dopamine Release in the Nucleus Accumbens Core Encodes the General Excitatory Components of Learning. The Journal of Neuroscience 44, e0120242024 (2024).

19. Sweis, B. M., Thomas, M. J. & Redish, A. D. Mice learn to avoid regret. PLoS Biol 16, (2018).

20. Frydman, C. & Camerer, C. Neural evidence of regret and its implications for investor behavior. in Review of Financial Studies vol. 29 3108–3139 (Oxford University Press, 2016).

21. Steiner, A. P. & Redish, A. D. Behavioral and neurophysiological correlates of regret in rat decision-making on a neuroeconomic task. Nat Neurosci (2014) doi:10.1038/nn.3740.

22. Coricelli, G., et al. Regret and its avoidance: A neuroimaging study of choice behavior. Nat Neurosci 8, 1255–1262 (2005).

23. Sweis, B. M., Redish, A. D. & Thomas, M. J. Prolonged abstinence from cocaine or morphine disrupts separable valuations during decision conflict. Nat Commun 9, (2018).

24. Sweis, B. M., Larson, E. B., David Redish, A. & Thomas, M. J. Altering gain of the infralimbic-to-accumbens shell circuit alters economically dissociable decision-making algorithms. Proc Natl Acad Sci U S A 115, E6347–E6355 (2018).

25. Durand-De Cuttoli, R., et al. Region-specific CREB function regulates distinct forms of regret associated with resilience versus susceptibility to chronic stress. Sci Adv (2022).

26. Sweis, B. M., et al. Sensitivity to “sunk costs” in mice, rats, and humans. Science (1979) 361, 178–181 (2018).

27. Durand-De Cuttoli, R., et al. Distinct forms of regret linked to resilience versus susceptibility to stress are regulated by region-specific CREB function in mice. Sci Adv (2022) doi:10.1126/sciadv.add5579.

28. Shansky, R. M. & Murphy, A. Z. Considering sex as a biological variable will require a global shift in science culture. Nature Neuroscience vol. 24 457–464 Preprint at 10.1038/s41593-021-00806-8 (2021).

29. Redish, A. D., et al. Sunk cost sensitivity during change-of-mind decisions is informed by both the spent and remaining costs. Commun Biol (2022) doi:10.1038/s42003-022-04235-6.

30. Mohebi, A., et al. Dissociable dopamine dynamics for learning and motivation. Nature 570, 65–70 (2019).

31. Klapoetke, N. C., et al. Independent optical excitation of distinct neural populations. Nat Methods (2014) doi:10.1038/nmeth.2836.

32. Gradinaru, V., et al. Molecular and Cellular Approaches for Diversifying and Extending Optogenetics. Cell 141, 154–165 (2010).

33. Lefevre, E. M., et al. Interruption of continuous opioid exposure exacerbates drug-evoked adaptations in the mesolimbic dopamine system. Neuropsychopharmacology 45, 1781–1792 (2020).

34. Nieuwenhuis, S., Forstmann, B. U. & Wagenmakers, E. J. Erroneous analyses of interactions in neuroscience: A problem of significance. Nature Neuroscience vol. 14 1105–1107 Preprint at 10.1038/nn.2886 (2011).

35. On the origin and early use of the term vicarious trial and error (VTE). https://psycnet.apa.org/fulltext/1958-02611-001.pdf.

36. Papale, A. E., Zielinski, M. C., Frank, L. M., Jadhav, S. P. & Redish, A. D. Interplay between Hippocampal Sharp-Wave-Ripple Events and Vicarious Trial and Error Behaviors in Decision Making. Neuron 92, 975–982 (2016).

37. Papale, A. E., Stott, J. J., Powell, N. J., Regier, P. S. & Redish, A. D. Interactions between deliberation and delay-discounting in rats. Cogn Affect Behav Neurosci 12, 513–526 (2012).

38. Redish, A. D. Vicarious trial and error. Nature Reviews Neuroscience 2016 17:3 17, 147–159 (2016).

39. Kay, K., et al. Constant Sub-second Cycling between Representations of Possible Futures in the Hippocampus. Cell 180, 552–567.e25 (2020).

40. Pan, W. X., Coddington, L. T. & Dudman, J. T. Dissociable contributions of phasic dopamine activity to reward and prediction. Cell Rep 36, (2021).

41. Coddington, L. T., Lindo, S. E. & Dudman, J. T. Mesolimbic dopamine adapts the rate of learning from action. Nature 614, 294–302 (2023).

42. Markowitz, J. E., et al. Spontaneous behaviour is structured by reinforcement without explicit reward. Nature 614, 108–117 (2023).

43. Freels, T. G., Gabriel, D. B. K., Lester, D. B. & Simon, N. W. Risky decision-making predicts dopamine release dynamics in nucleus accumbens shell. Neuropsychopharmacology 45, 266–275 (2020).

44. Le Heron, C., et al. Dopamine Modulates Dynamic Decision-Making during Foraging. Journal of Neuroscience 40, 5273–5282 (2020).

45. Amo, R., et al. A gradual temporal shift of dopamine responses mirrors the progression of temporal difference error in machine learning. Nat Neurosci 25, 1082–1092 (2022).

46. Maes, E. J. P., et al. Causal evidence supporting the proposal that dopamine transients function as temporal difference prediction errors. Nat Neurosci (2020) doi:10.1038/s41593-019-0574-1.

47. Lerner, T. N., Holloway, A. L. & Seiler, J. L. Dopamine, Updated: Reward Prediction Error and Beyond. Current Opinion in Neurobiology vol. 67 123–130 Preprint at 10.1016/j.conb.2020.10.012 (2021).

48. Gershman, S. J., et al. Explaining dopamine through prediction errors and beyond. Nature Neuroscience 2024 27, 1–11 (2024).

49. Kutlu, M. G., et al. Dopamine release in the nucleus accumbens core signals perceived saliency. Current Biology 31, 4748–4761.e8 (2021).

50. Jeong, H., et al. Mesolimbic dopamine release conveys causal associations. Science (1979) 378, (2022).

51. Starkweather, C. K., Babayan, B. M., Uchida, N. & Gershman, S. J. Dopamine reward prediction errors reflect hidden-state inference across time. Nat Neurosci (2017) doi:10.1038/nn.4520.

52. Hennig, J. A., et al. Emergence of belief-like representations through reinforcement learning. PLoS Comput Biol (2023) doi:10.1371/journal.pcbi.1011067.

53. Chiu, P. H., Lohrenz, T. M. & Montague, P. R. Smokers’ brains compute, but ignore, a fictive error signal in a sequential investment task. Nat Neurosci (2008) doi:10.1038/nn2067.

54. Coricelli, G., Dolan, R. J. & Sirigu, A. Brain, emotion and decision making: the paradigmatic example of regret. Trends in Cognitive Sciences Preprint at 10.1016/j.tics.2007.04.003 (2007).

55. Zeelenberg, M. & Pieters, R. Comparing service delivery to what might have Been: Behavioral responses to regret and disappointment. J Serv Res 2, 86–97 (1999).

56. Liu, C., et al. An inhibitory brainstem input to dopamine neurons encodes nicotine aversion. Neuron 110, 3018–3035.e7 (2022).

57. Stelly, C. E., et al. Pattern of dopamine signaling during aversive events predicts active avoidance learning. Proc Natl Acad Sci U S A (2019) doi:10.1073/pnas.1904249116.

58. Goedhoop, J. N., et al. Nucleus accumbens dopamine tracks aversive stimulus duration and prediction but not value or prediction error. Elife (2022) doi:10.7554/eLife.82711.

59. Iino, Y., et al. Dopamine D2 receptors in discrimination learning and spine enlargement. Nature 579, 555–560 (2020).

60. Tobler, P. N., Dickinson, A. & Schultz, W. Coding of Predicted Reward Omission by Dopamine Neurons in a Conditioned Inhibition Paradigm. Journal of Neuroscience (2003) doi:10.1523/jneurosci.23-32-10402.2003.

61. Schultz, W., Apicella, P. & Ljungberg, T. Responses of monkey dopamine neurons to reward and conditioned stimuli during successive steps of learning a delayed response task. Journal of Neuroscience 13, 900–913 (1993).

62. Saunders, B. T. & Robinson, T. E. Individual variation in the motivational properties of cocaine. Neuropsychopharmacology 36, 1668–1676 (2011).

63. Schmack, K., Bosc, M., Ott, T., Sturgill, J. F. & Kepecs, A. Striatal dopamine mediates hallucination-like perception in mice. Science (1979) (2021) doi:10.1126/science.abf4740.

64. Westbrook, A., et al. Dopamine promotes cognitive effort by biasing the benefits versus costs of cognitive work. Science (1979) 367, 1362–1366 (2020).

65. Tye, K. M., et al. Dopamine neurons modulate neural encoding and expression of depression-related behaviour. Nature (2013) doi:10.1038/nature11740.

66. Willmore, L., Cameron, C., Yang, J., Witten, I. B. & Falkner, A. L. Behavioural and dopaminergic signatures of resilience. Nature 611, 124–132 (2022).

67. Bäckman, C. M., et al. Characterization of a mouse strain expressing Cre recombinase from the 3′ untranslated region of the dopamine transporter locus. Genesis (United States) 44, 383–390 (2006).

68. Hart, W. E., Goldbaum, M., Côté, B., Kube, P. & Nelson, M. R. Measurement and classification of retinal vascular tortuosity. Int J Med Inform (1999) doi:10.1016/S1386-5056(98)00163-4.

69. Paxinos, G. & Franklin, K. B. J. The Mouse Brain in Stereotaxic Coordinates, 2nd edition. Academic Press Preprint at (2001).

